# Foxi1 regulates multipotent mucociliary progenitors and ionocyte specification through transcriptional and epigenetic mechanisms

**DOI:** 10.1101/2024.10.27.620464

**Authors:** Sarah Bowden, Magdalena Maria Brislinger-Engelhardt, Mona Hansen, Aisha Andricek, Africa Temporal-Plo, Damian Weber, Sandra Hägele, Fabian Lorenz, Tim Litwin, Clemens Kreutz, Peter Walentek

**Affiliations:** Internal Medicine IV, Medical Center - University of Freiburg, Hugstetter Strasse 55, 79106 Freiburg, Germany; CIBSS Centre for Integrative Biological Signalling Studies, University of Freiburg, Schänzlestrasse 18, 79104 Freiburg, Germany; IMPRS-IEM International Max Planck Research School of Immunobiology, Epigenetics and Metabolism, Max Planck Institute of Immunobiology and Epigenetics, Stübeweg 51, 79108 Freiburg, Germany; SGBM Spemann Graduate School for Biology and Medicine, University of Freiburg, Albertstrasse 19A, 79104 Freiburg, Germany; IMBI Institute of Medical Biometry and Statistics, Medical Center - University of Freiburg, Stefan-Meier-Strasse 26, 79104, Freiburg, Germany

**Author notes:** These authors contributed equally and are listed in alphabetical order (Sarah Bowden, Magdalena Maria Brislinger-Engelhardt, Mona Hansen).

**Keywords:** Xenopus epidermis, development, mucociliary, ionocytes, multipotent progenitors

## Abstract

Foxi1 is a master regulator of ionocytes (ISCs / INCs) across species and organs. Two subtypes of ISCs exist, and both α-and β-ISCs regulate pH-and ion-homeostasis in epithelia. Gain and loss of FOXI1 function are associated with human diseases, including Pendred syndrome, male infertility, renal acidosis and cancers. Foxi1 was predominantly studied in the context of ISC specification, however, reports indicate additional functions in early and ectodermal development. Here, we re-investigated the functions of Foxi1 in *Xenopus laevis* embryonic mucociliary epidermis development and found a novel function for Foxi1 in the generation of Notch-ligand expressing mucociliary multipotent progenitors (MPPs). We demonstrate that Foxi1 has multiple concentration-dependent functions: At low levels, Foxi1 maintains ectodermal competence in MPPs through transcriptional and epigenetic mechanisms, while at high levels, Foxi1 induces a multi-step process of ISC specification and differentiation in cooperation with Ubp1 and Dmrt2. We further describe how *foxi1* expression is affected through auto-and Notch-regulation, and how this developmental program affects mucociliary patterning. Together, we reveal novel functions for Foxi1 in *Xenopus* mucociliary epidermis formation, relevant to our understanding of vertebrate development and human disease.

## Introduction

The Forkhead-box transcription factor Foxi1 is a master regulator of ionocytes (ISCs) in the vertebrate lung, kidney, inner ear and epididymis (1, 2) as well as in the embryonic skin of aquatic species (e.g. zebrafish and *Xenopus* frogs) (3, 4). In all these tissues ISCs regulate ion homeostasis through the expression of transmembrane solute carriers and pH-regulators (e.g. vacuolar (v)H^+^ATPase encoded by *atp6* genes, Pendrin encoded by *slc26a4* and Anion exchanger 1 encoded by *slc4a1*)(3). In *Xenopus* embryos, an additional role for Foxi1 has been described during germ layer specification, where Foxi1 promotes epidermis formation by activating ectodermal gene expression while simultaneously counteracting vegetal mesendoderm-inducing factors (e.g. VegT) (5–7). However, how Foxi1 can have such profoundly different functions has not been elucidated so far.

The *Xenopus* embryonic epidermis is a popular model to study vertebrate mucociliary epithelia (8, 9). Mucociliary epithelia in the mammalian lung and the *Xenopus* epidermis serve as first line of defense against pathogens through mucociliary clearance (10). They are composed of secretory cells (e.g. goblet cells) that release mucus and anti-microbial peptides, and multi-ciliated cells (MCCs) that generate a fluid flow by means of directional cilia beating to remove pathogens (11). Mucociliary ISCs are important for efficient mucociliary clearance (12, 13), and Foxi1 dysregulation is linked to a range of human diseases affecting airway and kidney function as well as to causing deafness and cancers (2, 14–20). Hence, investigating how Foxi1 regulates diverse processes ranging from germ layer specification to cell type formation in the *Xenopus* mucociliary epidermis could reveal insights relevant to our understanding of vertebrate development as well as human diseases.

In this work, we elucidate novel concentration-dependent Foxi1 transcriptional and epigenetic functions in mucociliary development and ISC specification. In blastula and early gastrula stages, Foxi1 is expressed at lower levels and required to retain epidermal identity in a population of multipotent mucociliary progenitors (MPPs) residing in the deep layer of the prospective epidermis. MPPs are regulated by Foxi1 through transcriptional and epigenetic means, and are required for mucociliary cell type generation as well as Notch-mediated patterning. At high levels, Foxi1 then induces ISC specification in later gastrula and neurula stages. We further demonstrate that high Foxi1 expression in ISCs is achieved through auto-regulation, and that ISC differentiation is guided by additional transcription factors, Ubp1 and Dmrt2, which regulate α-and β-ISC subtype development.

## Results

### Foxi1 is expressed in multipotent mucociliary progenitors (MPPs)

Previous work in *Xenopus* has demonstrated that the maternally deposited transcription factor Foxi2 directly binds the *foxi1* promoter and activates its expression during zygotic genome activation (ZGA) (6). Recent work further demonstrated that Foxi2 cooperates with maternally deposited Sox3 in the early priming of genes for their activation during ZGA throughout the ectoderm (21), and that *foxi1* transcription is activated in the deep layer of the epidermal ectoderm (**Fig. 1A**). The deep ectodermal cell layer generates intercalating mucociliary cell types including ionocytes (ISCs), multiciliated cells (MCCs), and small secretory cells (SSCs) as well as basal cells (BCs) that serve as stem cells of the epidermis (**Fig. 1B**) (8, 9, 22).

**Figure 1:**
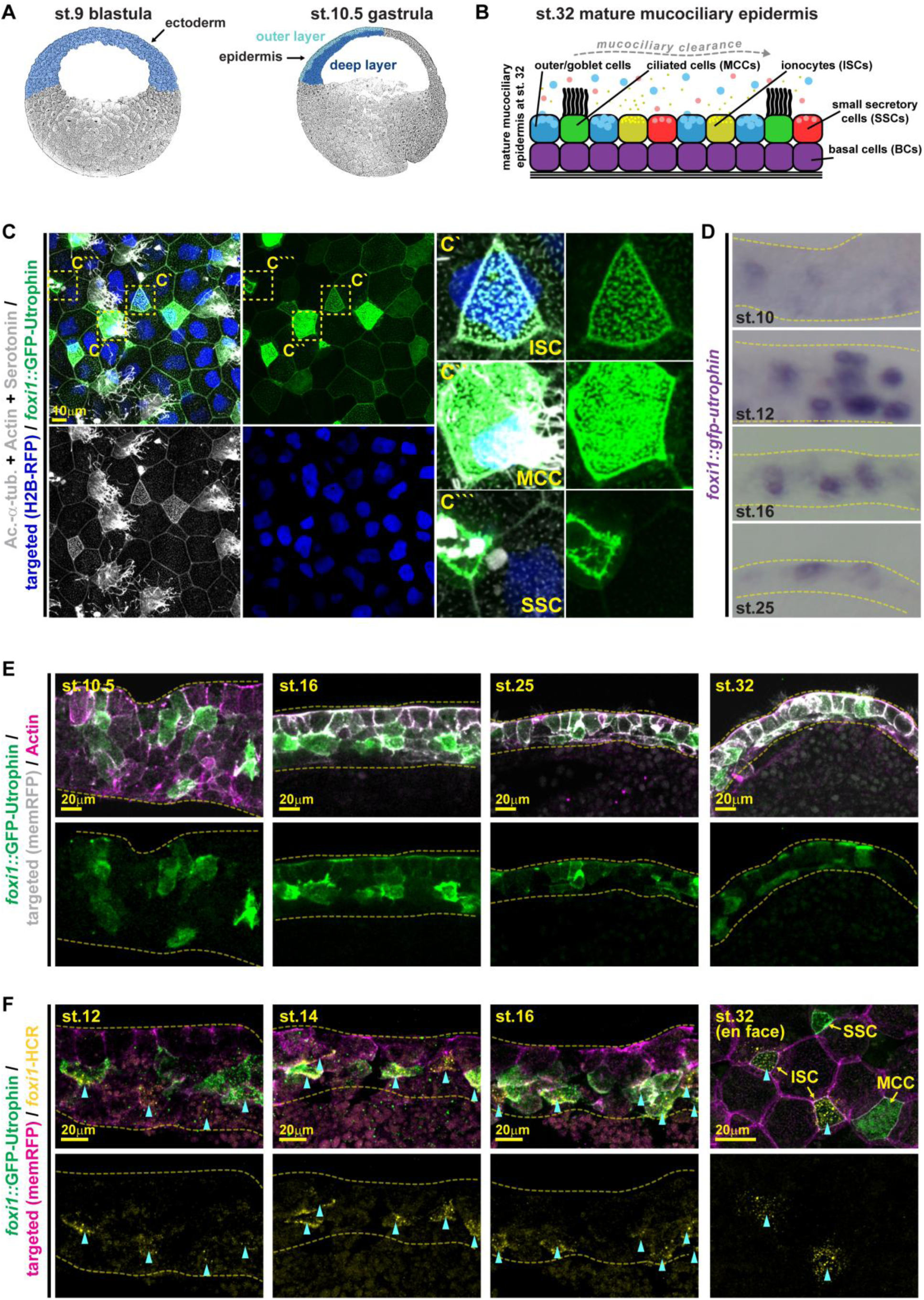
Foxi1 is transiently expressed in multipotent mucociliary progenitors (MPPs) **(A)** Schematic representations of a st. 9 blastula embryo (with prospective ectoderm expressing Foxi2 and Sox3 indicated in blue), of a st. 10.5 gastrula embryo (with outer layer epidermal cells indicated in light blue and the deep cell layer cells indicated in dark blue), and **(B)** of the mature (st. 32) mucociliary epidermis with basal cells (BCs) in purple, goblet cells (blue), ionocytes (ISCs) in yellow, ciliated cells (MCCs) in green, and small secretory cells (SSCs) in red. **(C-F)** Analysis of *foxi1::gfp-utrophin* reporter (green) injected embryos by IF **(C,E,F)** and WMISH against *gfp* **(D)**. **(C)** En face image, IF for Acetylated-α-tubulin (Ac.-α-tub, cilia, grey), F-actin (Actin, cell borders and morphology, grey), and serotonin in large vesicular granules (SSCs, grey) at st. 32. Targeted cells were identified by nuclear RFP expression (H2B-RFP, blue). Magnifications of intercalating GFP(+) cell types are shown in insets. Location of insets is indicated by dashed yellow boxes in left panels. n = 12 embryos. **(D)** Sections of epidermal locations from embryos depicted in Fig. S2C show *gfp* expressing cells in the epidermis at key stages of mucociliary development (st. 10, 16, 25, 32). st. 10 n = 19; st. 12 n = 16; st. 16 n = 14; st. 25 n = 14 embryos. **(E)** IF for *foxi1::gfp-utrophin* reporter (green) and F-actin (Actin, cell borders and morphology, magenta) at st. 10.5 - 32 on hemisected embryos. Apical up, basal down. Targeted cells were identified by membrane RFP expression (mRFP, grey). Additional stages shown in Fig. S3A. st. 10.5 n = 4; st. 16 n = 6; st. 25 n = 4; st. 32 n = 5 embryos. **(F)** HCR staining of endogenous *foxi1* transcripts (yellow) and IF for *foxi1::gfp-utrophin* reporter (green) at st. 12 - 16 on hemisected embryos and at st. 32 with en face view of the epithelium. Targeted cells were identified by membrane RFP expression (mRFP, magenta). Blue arrowheads indicate *foxi1*-HCR (+) cells. Stages st. 12 n = 5; st. 14 n = 4; st. 16 n = 3; st. 32 n = 4 embryos. Dashed lines demark the epidermis; apical up and basal down, in (D,E,F).

To re-evaluate Foxi1 functions in epidermis development, we first analyzed *foxi1* expression by whole mount *in situ* hybridization (WMISH). Shortly after ZGA, in early blastula/gastrula (st. 9/10) stages, *foxi1* was expressed at low levels in patches of the prospective ectoderm, which started to resolve at st. 12 with individual cells strongly increasing *foxi1* expression by st. 16, resulting in a salt-and-pepper pattern of individual cells by st. 32, representing individual ISCs (**Fig. S1A**). Hence, *foxi1* seemed to be transiently expressed in more epidermal cells than just in developing ISCs. We wondered if *foxi1* might be initially expressed at low levels in epidermal mucociliary multipotent progenitors (MPPs) in the deep epidermal layer. To test this, we generated a fluorescent reporter using the previously characterized *foxi1* promoter fragment harboring Foxi2 binding sites (6) driving the expression of GFP fused to the actin binding protein Utrophin for stable long-term labeling (*foxi1::gfp-utrophin*) (**Fig. S1B,C**). We injected embryos with *foxi1::gfp-utrophin* DNA and analyzed reporter activity at st. 32 by immunofluorescence (IF) and confocal microscopy. GFP signal was detected in ISCs, MCCs and SSCs, and even some goblet cells expressed GFP at low levels (**Fig. 1C**). In contrast, a Mcidas/Foxj1-regulated promoter construct driving mScarletI fluorescence (*α-tub::mscarletI*) was expressed predominantly in MCCs (**Fig. S2A,B**), as previously described (22, 23).

Next, we confirmed that temporal reporter expression dynamics resemble endogenous *foxi1* expression during epidermis development using WMISH (**Fig. 1D, S2C**) and GFP expression by IF (**Fig. 1E, S3A**). While plasmid injections lead to mosaic expression in the embryo, reporter-driven *gfp* transcripts were detected at st. 9 - 32, starting with non-epithelial low-level expression at st. 9/10, which increased by st. 12/16 in deep and superficial layer cells, and at st. 32 expression was found predominantly in epithelial layer cells (**Fig. 1D, S2C**). GFP-fluorescent cells were detected from st. 10 onwards, predominantly in deep layer cells, but also in some cells of the outer epithelial layer (**Fig. 1E, S3A**). During st. 12-16, an increasing number of cells became GFP(+), including intercalating differentiating cells (**Fig. 1E, S3A**). During st. 20 - 32, the number of GFP(+) cells decreased and fluorescent cells were progressively confined to the epithelial outer cell layer - however, basal positioned GFP(+) cells were detected even at st. 32 indicating that MPPs not differentiating into intercalating cell types become BCs (**Fig. 1E, S3A**). To further validate reporter specificity, we stained *foxi1::gfp-utrophin* injected embryos by fluorescent hybridization chain reaction (HCR) for endogenous *foxi1* transcripts. During cell fate specification stages (st. 12-16), most GFP(+) cells were also stained by *foxi1*-HCR, however, at st. 32, when the mucociliary epidermis is mature, only GFP(+) ISCs were still co-stained by *foxi1*-HCR in the epithelial layer, but not MCCs, SSCs or goblet cells (**Fig. 1F**).

Together these data support the conclusion that *foxi1* is initially expressed in MPPs during mucociliary epidermis development, and that *foxi1* is turned off in MCCs and SSCs during cell fate specification, while Foxi1 activity is maintained in ISCs.

### Foxi1 regulates genome accessibility of mucociliary genes in the epidermis

It was proposed that Foxi2 and Sox3 initially regulate broad epigenetic accessibility and gene expression in the ectoderm at ZGA, but that zygotic-expressed factors would be required to maintain accessibility and to drive gene expression in the epidermis during subsequent development (21). Besides its effects counteracting mesendoderm induction through transcriptional activation of ectodermal genes in early *Xenopus* embryos (7), Foxi1 has been shown to remain bound to condensed chromatin during mitosis, to remodel nucleosome structure and to alter the transcriptional ground state of cells in zebrafish embryos (24). This suggested that zygotic expression of *foxi1* could regulate both epigenetic state and transcriptional activity in epidermal MPPs.

To test if Foxi1 affects chromatin state and genomic accessibility in *Xenopus* epidermal development, we performed assays for transposase-accessible chromatin with sequencing (ATAC-seq) after morpholino-oligonucleotide (MO; 3 pmol) knockdown of *foxi1* (**Fig. S3B**). For these experiments, we used “animal cap”-derived organoids to specifically investigate chromatin state in pure epidermis tissue (25, 26). This analysis revealed a dramatic reduction in accessible chromatin regions (peaks) after loss of Foxi1 (control: 311,328, *foxi1* MO: 146,640) (**Fig.2A**,**B**). In Foxi1-depleted organoids, 53.5% of accessible regions (169,077 peaks) were lost, 45.1% were maintained (142,251 peaks), and 1.4% were gained (4,389 peaks) (**Fig. 2B**). Next, we investigated which transcription factor binding motifs were enriched in regions lost, maintained or gained after *foxi1* MO. We found that motifs for factors with known functions in *Xenopus* ectodermal development were enriched in regions that lost accessibility after *foxi1* knockdown: e.g. Tfap2a and Tfap2c, Hic1, Rbfox2, Zac1 that regulate neural and neural crest formation as well as Tp63, which regulates epidermal basal stem cells, and Pitx1, required for cement gland formation (**Fig. 2C**) (27–32). In contrast, regions that remained open were enriched for mesendodermal transcription factor motifs (e.g. Gata6, Tbxt, MyoD), and regions that gained in accessibility were enriched in pluripotency factors (e.g. Brn1, Oct4) (**Fig. 2C**) (33–37). Together, these data support a function for Foxi1 in retaining accessible chromatin state during epidermal development in *Xenopus*.

**Figure 2:**
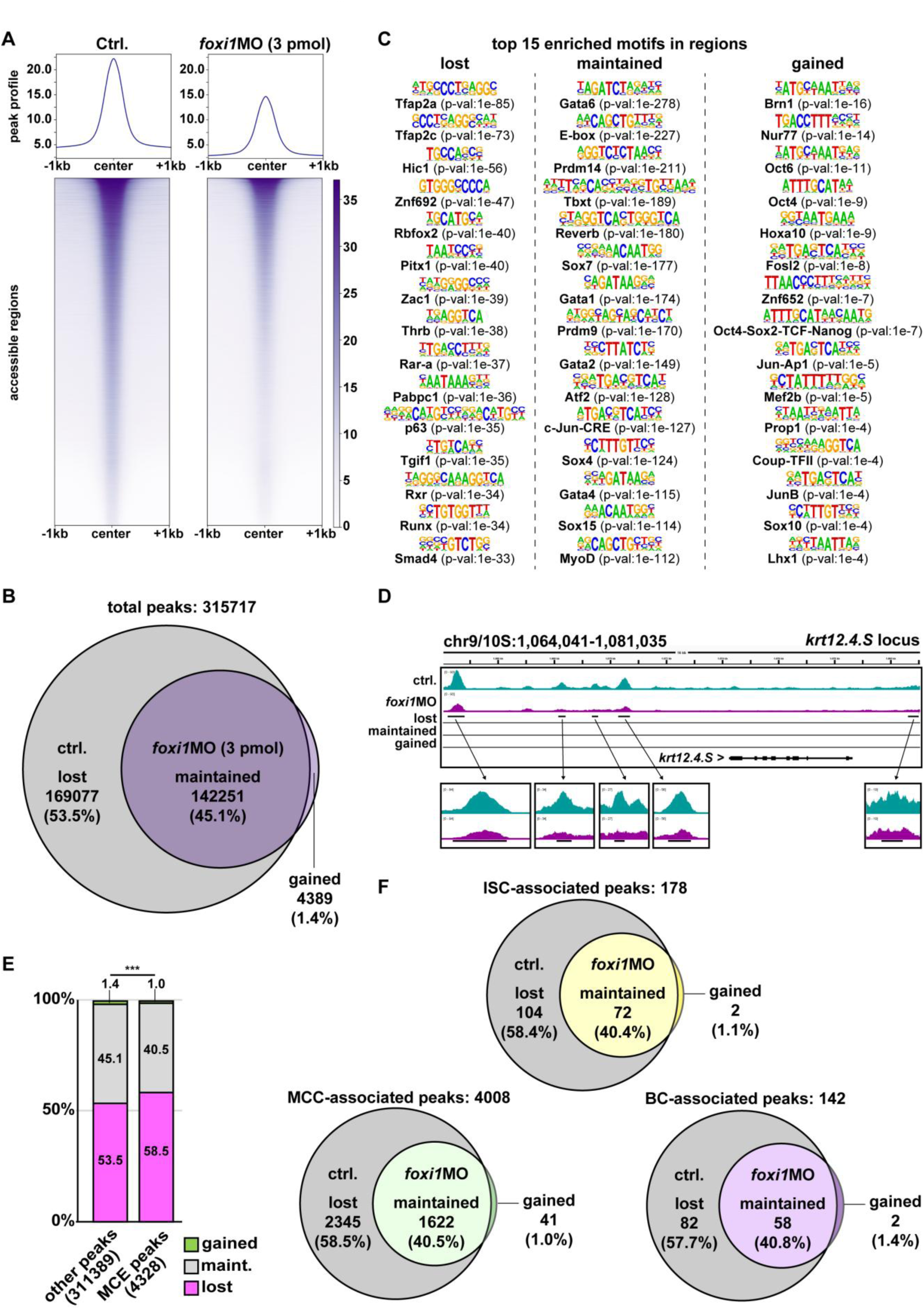
Foxi1 regulates chromatin state and mucociliary epidermal competence **(A)** Profiles of ATAC-Seq normalized accessibility around peak center ±1 kb in controls (ctrl.) and foxi1 morphant (*foxi1* MO, 3 pmol) organoids. n = 2 organoids per condition and replicate. 3 replicates. (**B)** Venn diagram of peaks present in uninjected organoids (grey) and *foxi1* MO-injected organoids (purple). (**C)** Top 15 transcription factor binding motifs predicted by HOMER in sets of peaks with lost, maintained or gained accessibility after *foxi1* MO. **(D)** Distribution of accessible regions around epidermal *krt12.4.* Lost, maintained and gained tracks as generated by MACS2 bdgdiff analysis and visualized in IGV. Turquoise track = control (ctrl.) and purple track = morphant (*foxi1* MO). **(E)** Enrichment of lost peaks in proximity of genes associated with mucociliary cell type development (MCE peaks). Green = gained; grey = maintained (maint.); magenta = lost. Chi² test: p < 0.001 = ***. **(F)** Venn diagrams of peaks present in uninjected organoids (grey) and *foxi1* MO-injected organoids (colored). ISC-associated peaks = yellow; MCC-associated peaks = green; BS-associated peaks = magenta.

Next, we wondered how loss of Foxi1 affects chromatin accessibility in regions harboring important genes for mucociliary epidermis development. First, we inspected a region around the *krt12.4* (epidermal keratin) gene on chromosome 9/10.S, which revealed strongly reduced accessibility and indicated a loss of epidermal competence (**Fig. 2D**) (7, 38). We further inspected genomic loci containing genes associated with mucociliary development (*dll1.L*), ISCs (*ubp1.L* and *dmrt2.S*), MCCs (*foxj1.L*) and BCs (*tp63.L*) (3, 39–41). In all cases, we found reduced accessibility (**Fig. S3C**). To investigate if the loss of Foxi1 particularly affected mucociliary genes as compared to other genes in the ectoderm, we investigated loci associated with published core-ISC,-MCC and-BC genes (40, 41). Strikingly, loss of accessibility in mucociliary gene-associated (MCE) peaks was significantly higher than in the rest of the genome (**Fig. 2E**), increasing from 53.7% lost non-MCE peaks to 58.4% lost ISC-associated peaks, to 58.5% lost MCC-associated peaks and to 57.7% lost BC-associated peaks (**Fig. 2F**).

In conclusion, Foxi1 regulates chromatin accessibility required for ectoderm and particularly for mucociliary cell type development. This provides an additional rational how Foxi1 could regulated mucociliary MPPs in early *Xenopus* epidermal development.

### Foxi1 acts in a concentration-dependent manner

Epidermal *foxi1* expression dynamics suggested that Foxi1 might act in a concentration-dependent manner: During early stages (st. 9-11), when MPPs are formed, *foxi1* expression is relatively low, while at stages when ISCs are specified and begin to differentiate (st. 12-16), *foxi1* expression dramatically increases in a subset of cells (**Fig. S1A**). To validate this observation using a quantitative approach and additional ISC markers, we used a previously defined a core-ISC gene set in the *X. laevis* (40) and investigated gene expression specifically in epidermal tissue using bulk RNA-sequencing (RNA-seq) on mucociliary organoids (25, 41, 42). Z-scores of normalized counts (TPM) of ISC transcripts were clustered to reveal dynamic co-expression (**Fig. 3A**). Five clusters clearly separated along developmental time, with cluster I being the only set of genes displaying strong expression during very early and late developmental stages, but not during cell fate specification stages (st. 10-16). Cluster II contained *foxi1*, the Notch ligand Delta-like 1 (*dll1*) and the cell cycle regulator *gadd45g*. Cluster III contained the pH-regulator Carbonic Anhydrase 12 (*ca12*; a pH regulator expressed in ISCs; (43)) and the transcription factor *ubp1*, which was shown to induce ectopic ISCs upon overexpression in the epidermis (3). Cluster IV contained multiple transcription factors, including *tfcp2l1*, required for ISC formation in the mouse kidney (44). Cluster V was dominated by solute carrier (e.g. *slc26a4*) and pH-regulator (*atp6*-subunits) expression during later differentiation of ISCs (st. 20–32). These data confirmed that *foxi1* expression levels were initially low (st. 9-11), and that they peaked at st. 12-14, shortly before the first functional ISC genes (e.g. *ubp1* and *ca12*) were expressed at st. 14-16 (**Fig. 3A**) (3, 43).

**Figure 3:**
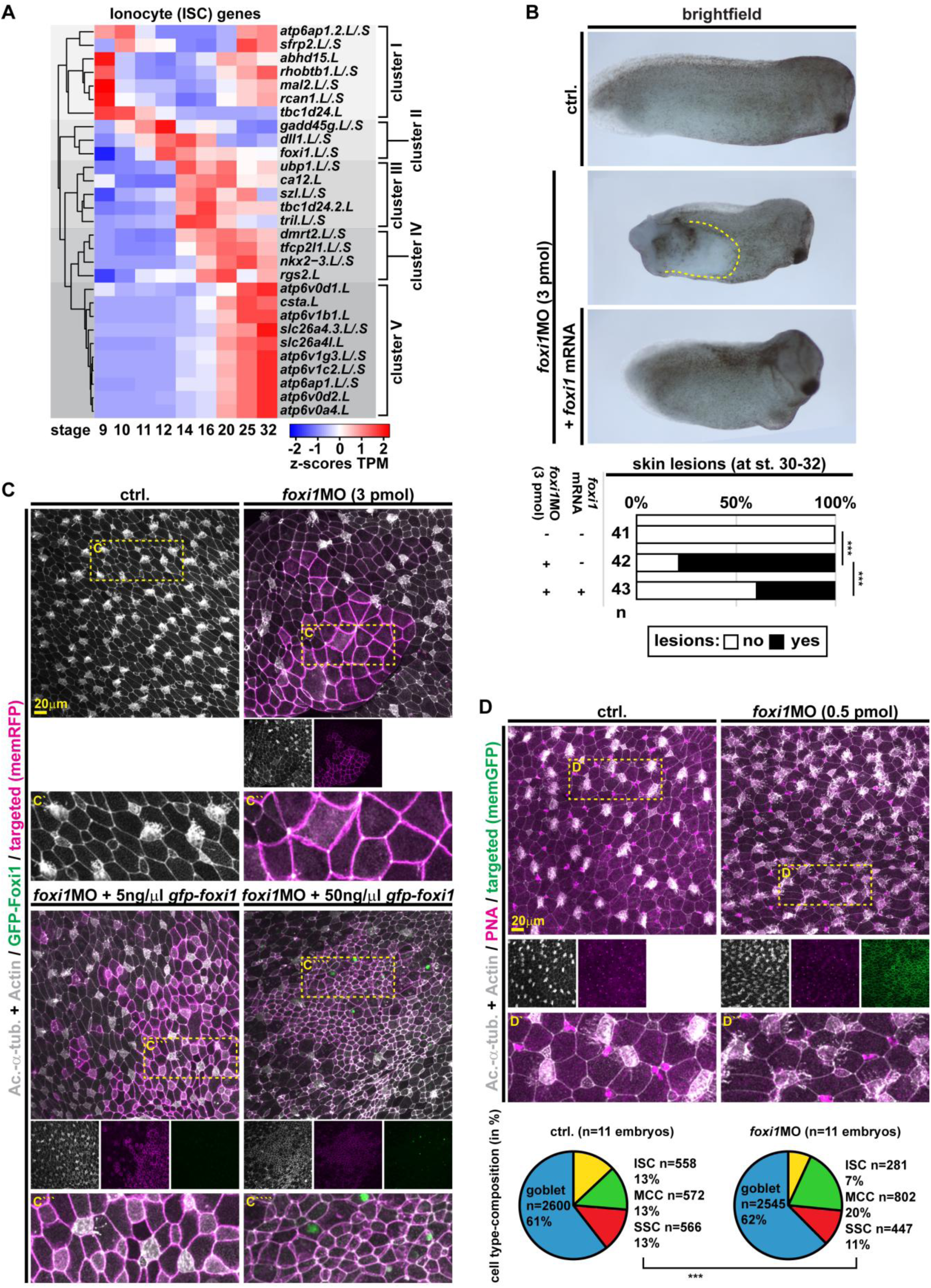
Foxi1 acts in a concentration-dependent manner **(A)** Temporal expression analysis of core ISC genes. Heatmap of line-normalized z-scores of TPMs (transcripts per million reads) derived from mRNA-seq of *Xenopus* mucociliary organoids over the course of development (st. 9 - 32). **(B)** Brightfield images and quantification of st. 30-32 embryos. Uninjected controls (ctrl.), *foxi1* morphants (*foxi1* MO; 3 pmol) and morphants co-injected with *foxi1* mRNA (50 ng/μl) are depicted anterior to the right, dorsal up. Skin lesions (dashed yellow outline) were quantified. Chi² test: p < 0.001 = ***. **(C,D)** Immunofluorescence confocal micrographs (IF) from control (ctrl.) and *foxi1* morphants (*foxi1* MO; concentrations are indicated) at st. 32 stained for Acetylated-α-tubulin (Ac.-α-tub, cilia, grey), F-actin (Actin, cell borders and morphology, grey). **(C)** Targeted cells were identified by membrane RFP expression (memRFP, magenta). Strongly targeted areas targeted by 3 pmol of *foxi1* MO fail to generate intercalating cell types, while co-injection of 5 and 50 ng/μl *gfp-foxi1* (green) rescues intercalated cell formation (identified by cilia staining or small apical surface of cells). Location of insets is indicated by dashed yellow box in upper panels. Ctrl. n = 9; *foxi1* MO n = 13; *foxi1* MO + 5 ng / μl = 7; *foxi1* MO + 5 ng / μl = 8. (**D**) To analyze cell type composition after low concentration of *foxi1* MO (1 pmol), mucus (PNA, magenta) was stained to reveal secretory cell types. Targeted cells were identified by membrane GFP expression (memGFP, green). Location of insets is indicated by dashed yellow box in upper panels. Quantification of cell type composition is depicted as pie-charts, goblet cells (blue), ISCs (yellow), MCCs (green) and SSCs (red). n embryos (above chart) and n quantified cells (in/left of chart). Chi² test: p < 0.001 = ***.

To test the hypothesis that high Foxi1 levels are required for ISC specification while lower Foxi1 levels are sufficient to retain epidermal identity, we first injected high *foxi1* MO doses (3 pmol; as used for the ATAC-seq experiments) targeted to the developing epidermis. This treatment induced delamination of cells without inducing apoptosis (TUNEL staining; (22)) in st. 9/10 embryos (**Fig. S4A,B**) and frequent formation of skin lesions at st. 32 (**Fig. 3B**) as previously described (5). The formation of skin lesions could be rescued by co-injection of *foxi1* mRNA (50 ng/μl), confirming MO specificity (**Fig. 3B**). Next, we performed IF on targeted regions of the epidermis at st. 32 to investigate mucociliary cell type formation and epidermal morphology. Cells receiving the highest doses of *foxi1* MO, marked by higher levels of co-injected lineage tracer (membrane RFP), did not form intercalating cell types including MCCs (revealed by acetylated-α-Tubulin staining in cilia) (**Fig. 3C**, **S4C**), as previously described (5). Next, we co-injected *foxi1* morphants with different concentrations of mRNA encoding GFP-tagged *foxi1* (*gfp-foxi1*) and analyzed rescue effects in the epidermis by IF. Low levels (5-15 ng/μl) of *gfp-foxi1* improved epithelial morphology and lead to the re-appearance of intercalating cells, including MCCs as well as cells with ambiguous morphology (**Fig. 3C**, **S4C**). High levels (50 ng/μl) of *gfp-foxi1* lead to a strong overproduction of intercalating cells with ISC-like morphology (**Fig. 3C**, **S4C**). However, a reliable assessment and quantification of cell type composition was limited by the severe morphological changes induced in morphants. Therefore, we tested the concentration-dependent rescue effects by quantitative PCR

(qPCR) at st. 10.5. We assessed a pan-epidermal marker (*krt12.4*) and a definitive-ISC marker (*ubp1*) on control and *foxi1* MO (3 pmol) injected samples as well as after co-injection of 5 – 50 ng/μl *gfp-foxi1* mRNA (**Fig. S4D**). This revealed a significant reduction in *krt12.4* in morphants, which was partially rescued by 15 ng/μl, but not 50 ng/μl of *gfp-foxi1*, while *ubp1* was significantly over-induced by 50 ng/μl of *gfp-foxi1*. Furthermore, we validated that different protein levels were induced at relevant stages (st. 12) using different *gfp-foxi1* mRNA concentrations by Western blot analysis (anti-GFP antibody) (**Fig. S4E**). While we could not assess endogenous Foxi1 protein level, collectively these data supported the hypothesis that lower levels of Foxi1 are sufficient to rescue epidermal identity and the formation of intercalating cell types from MPPs, while higher Foxi1 levels are required for ISC specification.

To confirm this, we injected low concentrations of *foxi1* MO (0.5 pmol) aiming at reducing only peak expression levels of Foxi1 without interfering with epidermis identity or MPPs, and analyzed cell type composition and morphology in the mature mucociliary epidermis at st. 32 by IF (25). This mild *foxi1* knockdown specifically reduced ISC formation without affecting epidermal identity as evidenced by the formation of other intercalating cell types (**Fig. 3D**): We observed little effects on secretory cells (SSCs and goblet cells) and while MCC ciliation was reduced as previously described in *X. tropicalis* (43), the overall number of MCCs was increased (**Fig. 3D**). This indicated that low concentrations of *foxi1* MO reduced Foxi1 levels enough to inhibit ISC specification (leading to supernumerary MCC specification), but not strong enough to interfere with ectoderm specification and MPP development.

This raised the question how MPPs can achieve high *foxi1* expression levels required for specification of ISC fate. One potential mechanism for conferring robust cell fate decisions is positive auto-regulation, and Foxi1 could activate its own expression using core Foxi motifs previously identified in the *foxi1* promoter (6, 45). To test this, we deleted a validated Foxi2 binding region (**Fig. S1B,C**) and analyzed reporter activity at st. 32, i.e. long after *foxi2* expression is terminated. This strongly decreased reporter activity (**Fig. S5A**), suggesting that core Foxi motifs are also used by Foxi1 to maintain its expression through auto-regulation. To verify that Foxi1 can also activate its own promoter without contributions from Foxi2, we injected *foxi1::gfp-utrophin* vegetally to target the prospective mesendoderm, which lacks maternally deposited *foxi2* (6). Analysis of reporter-only injected cells (marked by membrane RFP) in hemisected embryos at st. 11 showed no reporter activity in endodermal cells, while co-injection of *foxi1* mRNA led to ectopic activation of the reporter (**Fig. S5B**).

Together these data supported the conclusion that low levels of Foxi1 are sufficient for ectoderm identity and MPPs, that high Foxi1 levels are required for ISC specification, and that the *foxi1* expression increase in ISCs is achieved by positive auto-regulation (**Fig. 4A**).

**Figure 4:**
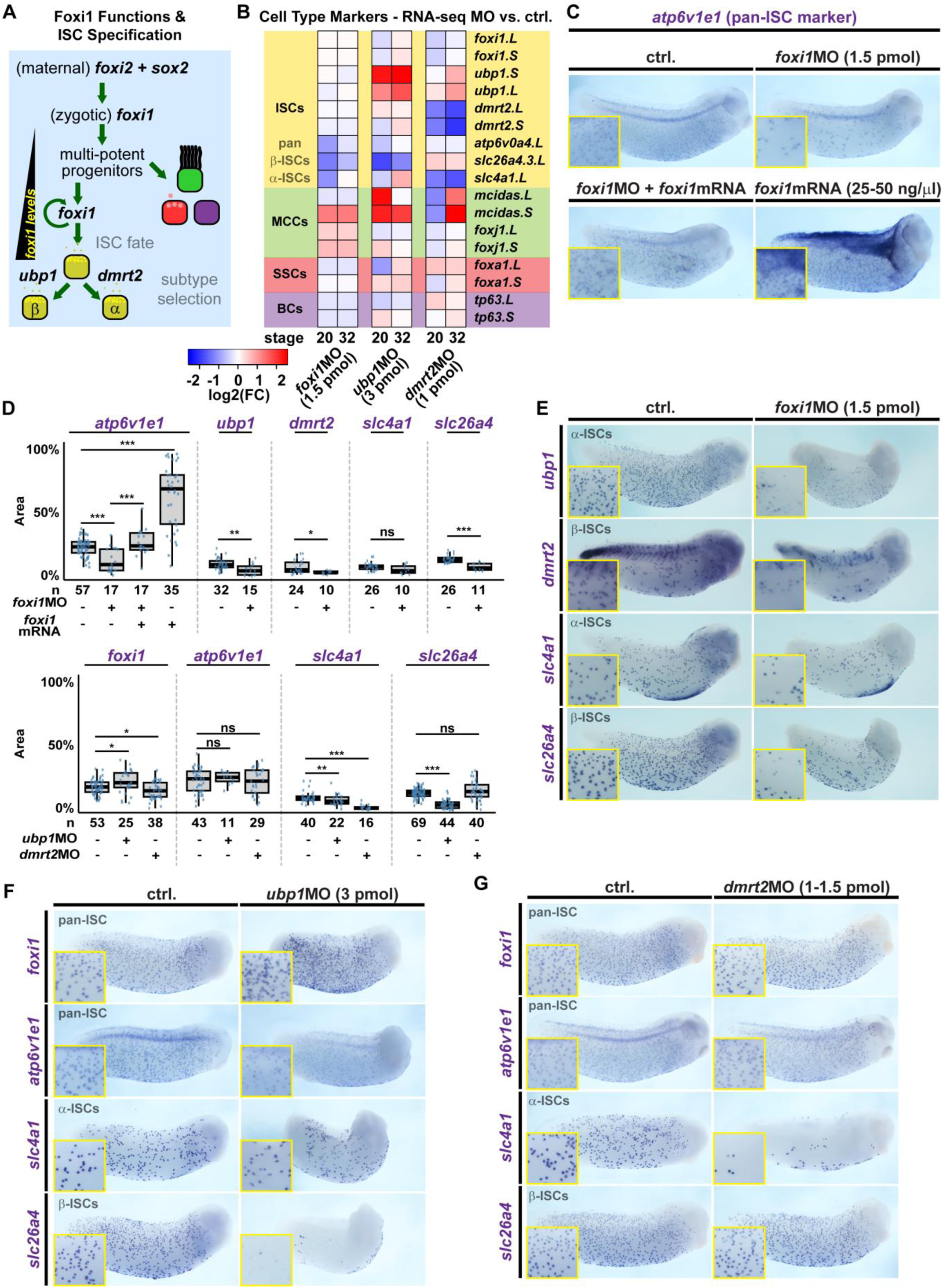
Foxi1, Ubp1 and Dmrt2 differentially regulate ionocyte development **(A)** Schematic representation of MPP and ISC specification in relation to transcription factor expression. **(B)** Gene expression analysis of mucociliary cell fate regulators after knockdown of Foxi1 (*foxi1* MO; 1.5 pmol), Ubp1 (*ubp1* MO; 3 pmol) and Dmrt2 (*dmrt2* MO; 1 pmol). Heatmap of log2-fold changes relative to control samples derived from DEseq2 analysis of mRNA-seq on *Xenopus* mucociliary organoids during ISC differentiation (st. 20) and in mature epidermis stage (st. 32). **(C-G)** Knockdown of ISC transcription factors (*foxi1* MO, 1.5 pmol; *ubp1* MO, 3 pmol; *dmrt2* MO, 1 pmol) and analysis of effects by WMISH at st. 29 - 32 against *atp6v1e1* and *foxi1* (pan-ISC markers), *ubp1* and *slc25a4/pendrin* (β-ISC markets), and *dmrt2* and *slc4a1/ae1* (α-ISC markers). Representative images, embryos depicted anterior to the right, dorsal up, insets show magnification of epidermis in **(C,E,F,G)** and quantification of results in **(D)**. n = number of embryos analyzed per condition. **(C)** Co-injection of 25-50 ng/μl of *foxi1* mRNA rescues expression of pan-ISC marker *atp6v1e1* in *foxi1* morphants (1.5 pmol) and *foxi1* overexpression induces broad *atp6v1e1* expression in the epidermis. In **(D)**, marker positive area within a standardized region of embryo was analyzed and is depicted as % of analyzed area. Wilcoxon Rank Sum test: p > 0.05 = ns; p < 0.05 = *; p < 0.01 = **; p < 0.001 = ***. **(D,E)** Foxi1 knockdown (*foxi1* MO, 1.5 pmol) leads to loss of marker expression in both ISC subtypes. **(D,F)** Ubp1 knockdown (*ubp1* MO, 3 pmol) leads to loss of β-ISC marker. **(D,G)** Dmrt2 knockdown (*dmrt2* MO, 1 pmol) leads to loss of α-ISC marker.

### Specification and differentiation of ISCs is a multi-step process

Besides concentration-dependent effects of transcription factors, an important route to modulate transcription factor functions during development is the cooperation with other transcription factors (46). Therefore, we next investigated how other transcription factors contained in the core-ISC gene set regulate ISC specification (**Fig. 3A**). Among the first definitive ISC genes expressed during mucociliary epidermis development were the transcription factors Ubp1 (Cluster III) and Dmrt2 (Cluster IV). Two ISC subtypes exist, α-ISCs that differentially express AE1 (encoded by *slc4a1*) and β-ISCs that express Pendrin (encoded by *slc26a4*) (3). Additionally, both ISC subtypes express vH+-ATPase (encoded by *atp6*-subunits)(3, 47). Ubp1 was previously shown to induce β-ISCs in *Xenopus*, and Dmrt2 was recently shown to be required for α-ISC formation in the mouse kidney (3, 48).

We used single-cell RNA-seq (scRNA-seq) data from at late *Xenopus laevis* tail stages containing mature ISCs (49) to investigate α-and β-ISC gene profiles. These data revealed high levels of *foxi1* and *atp6*-subunit expression in both ISC subtypes, while *ubp1* was enriched and *slc26a4* was specifically expressed in β-ISCs (**Fig. S5C**). In contrast, *slc4a1* as well as *dmrt2* were expressed only in α-ISCs (**Fig. S5C**). To reveal ISC-subtype specific functions, we tested how *foxi1*, *ubp1* and *dmrt2* contribute to ISC-subtype formation by RNA-seq on mucociliary organoids during differentiation (st. 20) and when the epidermis is mature (st. 32) (**Fig. 4B**). Knockdown of Foxi1 (1.5 pmol *foxi1* MO) should reduce expression of all ISC genes, while knockdown of Ubp1 (3 pmol *ubp1* MO) and Dmrt2 (1 pmol *dmrt2* MO) should yield differential effects on the subtype-specific ISC markers (*slc4a1* – α-ISCs; *slc26a4* – β-ISCs) (**Fig. S3B**). RNA-seq on *foxi1* morphant organoids revealed a reduction of ISC differentiation markers *atp6*, *slc26a4* and *slc4a1* (**Fig. 4B**), and the expression of core-ISC genes was reduced across all clusters (**Fig. S6A**). Furthermore, the MCC genes *mcidas* and *foxj1* were upregulated, in line with our finding that MCCs are over-produce when ISC specification was blocked (**Fig. 3D**). In contrast to *foxi1* MO injections, knockdown of *ubp1* caused strong downregulation of *slc26a4*, while *atp6* was only transiently reduced and *slc4a1* was elevated at st. 32 (**Fig. 4B**). A reduction of ISC-gene expression was predominantly found in cluster I and V genes, while genes in cluster III were upregulated (**Fig. S6B**). This indicated a specific loss of β-ISCs as well as a change in differentiation dynamics. Knockdown of *dmrt2* caused strong downregulation of *slc4a1*, while *atp6* remained unchanged and *slc26a4* was elevated (**Fig. 4B**). Across core-ISC genes, *dmrt2* MO only led to a strong downregulation of *dmrt2* expression, while most genes across all clusters were only transiently reduced (**Fig. S6C**). This indicated a specific loss of α-ISCs. Furthermore, differentiation dynamics appeared to be dysregulated after *dmrt2* MO, and similar to *foxi1* as well as *ubp1* MOs, an upregulation of *mcidas* was observed (**Fig. 4B, S6C**).

To validate the findings from manipulating organoids, we knocked down each factor and analyzed ISC marker expression in the mature epidermis at st. 30-32 by WMISH. *foxi1* (MO, 1.5 pmol) knockdown strongly reduced the pan-ISC marker *atp6v1e1* expression, which could be rescued by co-injection of *foxi1* mRNA (**Fig. 4C,D**). Furthermore, 1.5 pmol of *foxi1* MO reduced *ubp1*, *dmrt2*, *slc24a6* and *slc4a1* expression, in line with the RNA-seq data (**Fig. 4B,D,E**), indicating a loss of both ISC subtypes. In contrast, knockdown of *ubp1* specifically reduced expression of the β-ISC marker *slc26a4*, while *dmrt2* loss lead to inhibition of α-ISC-specific *slc4a1* expression (**Fig. 4D,F,G**). Importantly, *ubp1* and *dmrt2* MOs did not abolish pan-ISC expressed *foxi1* and *atp6v1e1* (**Fig. 4D,F,G**). MO-specific effects were confirmed in rescue experiments, and ISC-specific mRNA effects were confirmed by overexpression of *ubp1* and *dmrt2* causing supernumerary induction of ISC subtypes (**FIG S5D,E**).

Taken together, these data support the conclusion that Foxi1 determines ISC fate commitment, while Ubp1 and Dmrt2 cooperate with Foxi1 during ISC-subtype differentiation in a multi-step process of ISC development (**Fig. 4A**).

### MPPs and ISC transcription factors regulate Notch signaling during cell fate specification

Previous studies indicated that inhibition of Foxi1 at a level that prevents specification of ISCs but not MCCs negatively affected normal MCC ciliation (43). In line with this observation, our data indicated that while MCC numbers were increased after *foxi1* MO (0.5 pmol), ciliation was reduced in many MCCs (inset in **Fig. 3D**). Previously, we have shown that elevated Notch levels presented to MCCs after fate commitment interfered with normal ciliation and differentiation (22). Hence, we hypothesized that a dysregulation of Notch signaling could cause the MCC ciliation phenotype. Notch signaling is required for mucociliary cell fate decision across mucociliary epithelia, and in the *Xenopus* epidermis, ISCs as well as MCCs were shown to be inhibited by Notch activity (3, 39). The Notch ligand *dll1* is expressed during epidermis development, and its expression was assigned to ISCs by Quigley and colleagues, similar to Foxi(+) cells in the zebrafish skin and mammalian kidney (4, 40, 50). However, another study observed *dll1* expression overlapping with different cell markers during patterning stages in the *Xenopus* epidermis (51). Our temporal expression analysis indicated very early *foxi1* and *dll1* expression in Cluster II, likely representing MPPs and early ISC differentiation stages (**Fig. 3A**), in line with both published observations. Therefore, Foxi1 manipulations could dysregulate MPPs expressing *dll1*, and thereby alter Notch signaling during patterning in the epidermis.

To investigate how MPPs, Notch signaling and core-ISC genes are regulated in the mucociliary epidermis, we manipulated Notch and cell fates in organoids and investigated gene expression at st. 10.5, 16, 25, and 32. As previously described (3), increased Notch signaling (Notch intracellular domain (*nicd*) mRNA injections) inhibited core ISC gene expression, while inhibition of Notch signaling (injection of dominant-negative suppressor of hairless/RBPJ (*suh-dbm*) mRNA) promoted core ISC gene expression (**Fig. S7A,B**). Inhibition of Notch signaling in combination with blocking MCCs (by co-injection of *dominant-negative mcidas* (*dn-mcidas*) mRNA (52)) further increased core ISC gene expression, reflecting stronger overproduction of ISCs. However, expression of *tfcp2l1*, *atp6v0d1.L* and *csta.L* were reduced in these conditions suggesting that they might be not be specifically expressed in ISCs (**Fig. S7C**). Notch repression of *foxi1* by *nicd* was substantial but not statistically significant (**Fig. S7A**), likely reflecting a Notch-independent regulation of Foxi1 in MPPs that is retained when the specification of intercalating cell types is inhibited.

Feedback regulation of *dll1* by Notch signaling was suggested in *Xenopus* epidermis development (39). RNA-seq analysis of *dll1* expression after Notch manipulations confirmed that gain of Notch signaling suppresses *dll1*, while blocking Notch increases and prolongs *dll1* expression (**Fig. S7A,B**). Interestingly, blocking MCC formation in Notch-inhibited organoids further increased and prolonged *dll1* expression, indicating that MCC fate specification inhibits *dll1* expression when MPPs adopt this cell fate (**Fig. S1B,C**). To address if *dll1* (and *dlc*; Brislinger-Engelhardt et al, 2023) expression is part of the differentiation program across multiple mucociliary cell types or specific to MPPs and early ISCs, we tested whether master transcription factors inducing cell fates of the mucociliary epidermis were able to induce Notch ligands prematurely. To induce the different cell types, we overexpressed *foxi1* for MPPs/ISCs, *mcidas* and *foxj1* for MCCs, *foxa1* for SSCs, and for BCs *ΔN-tp63* (Tp63 isoform that regulates mucociliary BCs (41)). Only *foxi1* robustly induced *dll1* and *dlc* (**Fig. 5A, S7D**), and conversely, depletion of Foxi1 (3 pmol MO) prevented *dll1* expression during cell fate specification stages (**Fig. 5B**).

**Figure 5:**
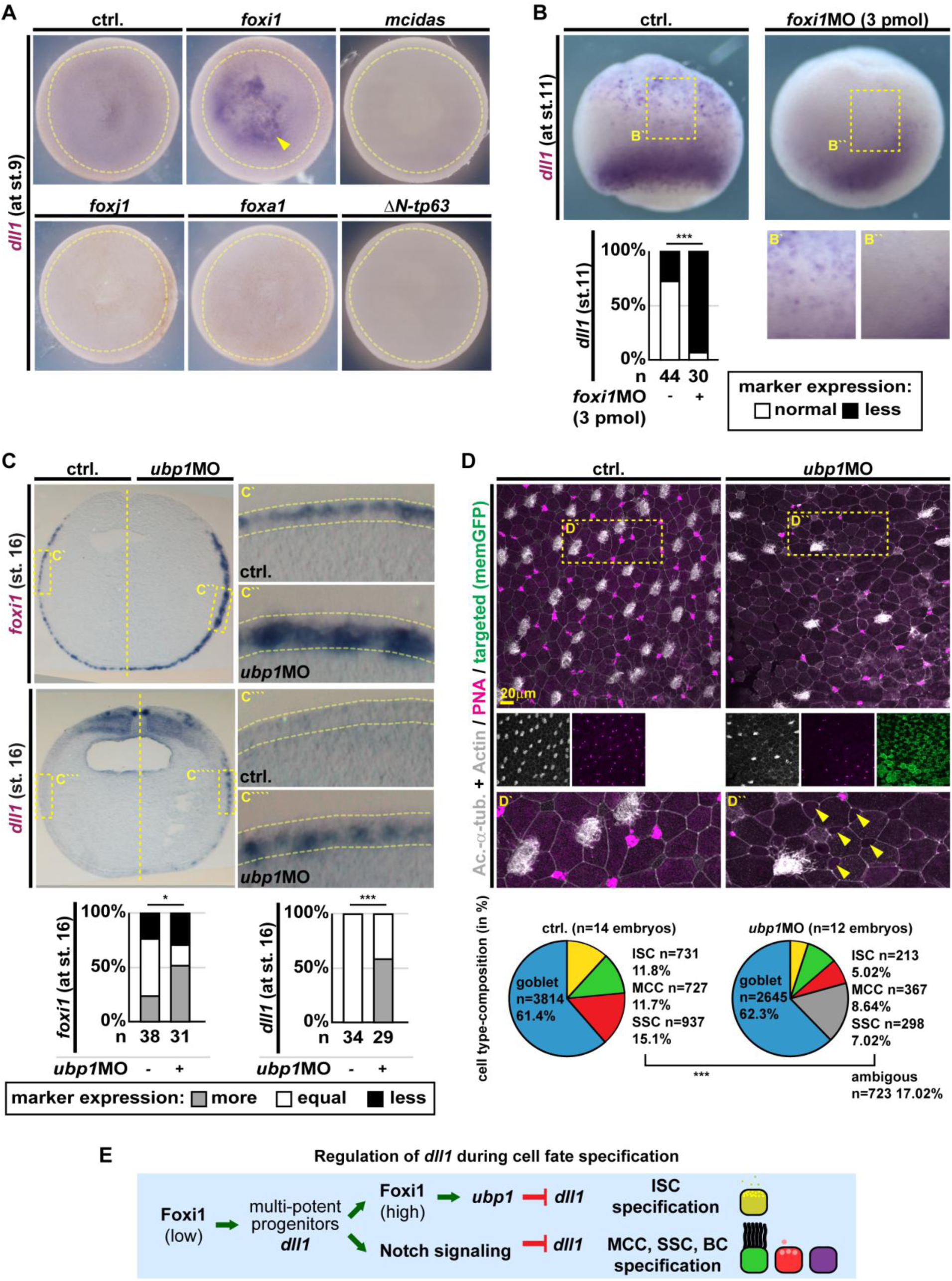
Foxi1 induces and Ubp1 terminates Notch ligand expression **(A,B,C)** Manipulation of mucociliary cell fate transcription factors *foxi1* and *ubp1* (ISCs/MPPs), *mcidas* and *foxj1* (MCCs), *foxa1* (SSCs) and *ΔN-tp63* (BCs) and analysis of effects by WMISH at st. 9 (animal view, ectoderm outlined in yellow) **(A)**, st. 11 (ventro-lateral view, blastopore down, animal regions up) **(B)** and st. 16 section of unilateral injected embryos; dorsal up, ventral down) **(C)** against *dll1* (Notch ligand) and *foxi1* (MPP/ISC marker). **(A)** Representative images of control (ctrl.) and manipulated embryos (animal views) after mRNA overexpression of transcription factors to test premature induction of *dll1*. Quantification of results and effects on *dlc* are shown in Fig. S7D. Embryos were scored as induced or non-induced expression. Yellow arrowhead indicates induced expression. **(B)** Representative images of control (ctrl.) and *foxi1* morphants (*foxi1* MO, 3 pmol) to test effects on *dll1* expression. Quantification of results is shown in lower panel. Insets show epidermal area magnification. Locations of insets are indicated by dashed yellow box in upper panels. Embryos were scored as normal or reduced (less) expression of *dll1*. Chi² test: p < 0.001 = ***. **(C)** Representative images of section embryos after unilateral knockdown of *ubp1* (*ubp1* MO, 3 pmol). Expression of markers was scored as more, less or equal to uninjected control (ctrl.) side. Locations of insets are indicated by dashed yellow box in left panels. Epidermis is outlined in magnified images, apical up and basal down. Chi² test: p < 0.05 = *; p < 0.001 = ***. **(D)** IF of control (ctrl.) and *ubp1* morphants (*ubp1* MO, 3 pmol) at st. 32 stained for Acetylated-α-tubulin (Ac.-α-tub, cilia, grey), F-actin (Actin, cell borders and morphology, grey), and mucus (PNA, magenta). Targeted cells were identified by membrane GFP expression (memGFP, green). Location of insets is indicated by dashed yellow box in upper panels. Ambigous cells are indicated by yellow arrowheads. Quantification of cell type composition is depicted as pie-charts, goblet cells (blue), ISCs (yellow), MCCs (green) and SSCs (red). Ambigous cells are depicted in grey. n embryos (above chart) and n quantified cells (in/left of chart). Chi² test: p < 0.001 = ***, not including ambiguous cells. **(D)** Schematic summary of Dll1 regulation during mucociliary development.

These results suggested that *dll1* is expressed in *foxi1*(+) MPPs and terminated by Notch signaling and cell fate induction of MCCs, SSCs and BCs.

In differentiating and mature ISCs, which maintain *foxi1* expression, *dll1* expression should be also maintained. Nevertheless, RNA-seq data indicate that *dll1* expression is not maintained at high levels in the mature mucociliary epidermis (**Fig. 3A**). This raised the question how *dll1* expression is terminated during ISC differentiation. To address that, we investigated published developmental *X. tropicalis* scRNA-seq data (53) and visualized enrichment for key mucociliary cell fate regulators and *dll1* across cell types and developmental stages. This confirmed that *dll1* is enriched early in the ISC lineage, and that *dll1* expression is lost once *ubp1* is expressed (**Fig. S7E**). Furthermore, our RNA-seq data on manipulated organoids confirmed an upregulation of *dll1* when ISC differentiation was inhibited, and these effects were most pronounced after *ubp1* MO (**Fig. S6A-C**). To validate that Ubp1 terminates *dll1* expression in embryos, we knocked down *ubp1* and analyzed embryos at the end of cell fate specification (st. 16) by WMISH. *foxi1* was maintained and *dll1* expression was prolonged on the *ubp1* MO injected side (**Fig. 5C**). Analysis of cell type composition at st. 32 by IF in *ubp1* morphants further revealed reduced MCC and SSC numbers as well as appearance of intercalating cells with ambiguous morphology, likely representing incompletely differentiated ISCs (**Fig. 5D**). These results revealed that manipulating *foxi1* and ISC differentiation leads to dysregulated Notch dynamics during mucociliary development. This explains why we and others have observed MCC ciliation defects after Foxi1 manipulations.

Together, these data suggest that Notch ligands are expressed by *foxi1*(+) MPPs during mucociliary patterning, and that Ubp1 terminates *dll1* expression in the ISC lineage.

Furthermore, induction of MCC, SSC and BCs also terminates Notch ligand expression in other differentiating mucociliary cell types. This provides a mechanism to regulated Notch signaling during mucociliary patterning that is tuned by both feedback-regulation (Notch) as well as fate commitment induced by mucociliary transcription factor networks (**Fig. 5E**).

## Discussion

This work revealed that Foxi1 regulates multiple crucial steps during *Xenopus* mucociliary epidermis development through transcriptional and epigenetic mechanisms. Initially, low-level *foxi1* expression is activated in the prospective ectoderm by maternally deposited *foxi2* and *sox2* (6, 21). This low-level expression is required to regulate chromatin accessibility for pro-ectodermal transcription factors (e.g. Tfap2a/c), mucociliary regulators (e.g. Tp63) as well as mediators of thyroid hormone, retinoic acid and TGFβ signaling (e.g. Thrb, Rar-a, Smad4) that were described to regulate ectodermal development (22, 41, 51, 54, 55). Since baseline *foxi1* expression is Foxi2-/Sox2-but not Notch-dependent, Notch over-activation does not alter ectodermal identity in line with previous reports (6, 21, 39). Hence, Foxi1 could maintain epidermis developmental potential in deep layer cells after the activities of maternal factors Foxi2 and Sox2 are reduced during gastrula stages (21).

Furthermore, our data support previous findings that loss of Foxi1 leads to acquisition of mesendodermal fates, as loci enriched for pro-mesendodermal transcription factors (e.g. Gata6, Tbxt, MyoD) remain accessible in the absence of Foxi1 (33–35). It is attractive to speculate that this is possible, because multiple transcription factors enriched in the maintained fraction of peaks (e.g. Gata and Sox family members) are known factors with pioneer activity (56, 57).

After regulating MPPs during early development, Foxi1 levels then increase through auto-regulation, and high levels of Foxi1 induce ISC fate in cooperation with Ubp1 and Dmrt2, in line with the known role of Foxi1 as master transcription factor for ionocytes across vertebrate tissues (1). ISC fate, subtype selection and differentiation is a multi-step process. In the *Xenopus* epidermis, Ubp1 initially terminates Notch ligand expression in differentiating α-and β-ISCs. Then, Ubp1 drives differentiation of β-ISCs, while Dmrt2 drives α-ISC differentiation. This highlights the importance of transcription factor cooperativity in cell fate decisions, in addition to Foxi1 concentration-dependent effects. Interestingly, Dmrt2 has recently been shown to be required for α-ISCs in the mouse kidney; however, not Ubp1, but Tfcp2l1 is employed in mammalian kidney β-ISCs (3, 44, 48). Differential use of these grainyhead-like transcription factors could explain why Dll1 expression is terminated in *Xenopus* epidermal ISCs, but remains active in mammalian kidney ISCs (also called INCs) (44, 50). Similar to Ubp1, our results suggest that fate acquisition of other mucociliary epidermal cell types terminates Dll1/Dlc ligand expression in MPPs. Together, this system provides a robust Notch feedback-regulated developmental program for mucociliary epidermis development (58), with Foxi1 as a central player that acts through transcriptional as well as epigenetic mechanisms, and that affects cell fate specification directly in ISCs as well as indirectly through dysregulating Notch ligand expression in MPPs. However, one limitation at this point is that we have not found a specific marker exclusively expressed in MPPs, which would be necessary to investigate MPP-specific behavior and gene expression further.

Finally, in ISCs of the mammalian airway mucociliary epithelium, Foxi1 also regulates the expression of *cystic fibrosis transmembrane conductance regulator* (*CFTR*), and mutations in *FOXI1* and its transcriptional target solute carriers cause Pendred syndrome and hearing loss, male infertility, and distal renal tubular acidosis (2, 13–15). In contrast, Foxi1 overexpression is found in cancer subtypes, e.g. in chromophore renal cell carcinoma and in pulmonary large cell carcinoma, but how Foxi1 overactivation can lead to cancerous transformations remains unresolved (17–20). Hence, our finding that Foxi1 drives an MPP state during mucociliary epidermis development could serve as a starting point to better understand the role of Foxi1 in these cancers.

## Acknowledgments

We thank: S. Schefold for expert technical help; L. Kodjabachian and team, T. Manke and W. Deboutte, T. Kwon for support and discussions; P. Klein for ATAC protocol; Xenbase, EXRC for Xenopus resources; Light Imaging Center Freiburg, BiMiC and Aqua Core for microscope/animal resources; B. Grüning and the Freiburg Galaxy Team for bioinformatics platform and support. This study was supported by the Deutsche Forschungsgemeinschaft (DFG) under the Emmy Noether and Heisenberg Programmes (grant WA3365/2-1 and WA3365/5-1), by DFG SFB1453 NephGen (Project ID 431984000), by DFG/ANR grant WA3365/4-1, and by the NHLBI through a Pathway to Independence Award (K99HL127275) to PW; and under Germany’s Excellence Strategy (CIBSS – EXC-2189 – Project ID 390939984) to PW and CK.

## Author contribution

SB: epigenetics; MMBE: cell fates and Notch; MOH: reporter studies; AA, ATP, DW, SH, PW: experimental support; SB, MMBE, MOH, PW: experimental design, planning, analysis and interpretation of data; FL, TL, CK: crucial discussion and mathematical modeling; PW, SB: bioinformatics. PW: study design and supervision, coordinating collaborative work, manuscript preparation with input from all authors. SB, MMBE, MOH contributed equally and can list themselves as first co-first authors.

## Material and Methods

### Animal experiments

Wild-type *Xenopus laevis* were obtained from the European Xenopus Resource Centre (EXRC) at University of Portsmouth, School of Biological Sciences, UK, or Xenopus 1, USA. Frog maintenance and care was conducted according to standard procedures in the AquaCore facility, University Freiburg, Medical Center (RI_00544) and based on recommendations provided by the international Xenopus community resource centers NXR (RRID:SCR_013731) and EXRC as well as by Xenbase (http://www.xenbase.org/, RRID:SCR_003280)(59). This work was done in compliance with German animal protection laws and was approved under Registrier-Nr. G-18/76 and G-22/43 by the state of Baden-Württemberg.

### Data availability

NGS datasets are available via NCBI GEO, ATAC-seq datasets (GSE280790), mRNA-seq datasets that were generated in previous studies (GSE130448, GSE215373, GSE215419, GSE262944)(41, 42), mRNA-seq datasets that were generated for this study (GSE299718). Imaging and quantification data are available to the scientific community upon request to.

### Manipulation of *Xenopus* Embryos

*X. laevis* eggs were collected and in vitro-fertilized, then cultured and microinjected by standard procedures (60, 61). Embryos were injected with Morpholino oligonucleotides (MOs, Gene Tools), mRNAs or plasmid DNA at two-cell to eight-cell stage using a PicoSpritzer setup in 1/3x Modified Frog Ringer’s solution (MR) with 2.5% Ficoll PM 400 (GE Healthcare, #17-0300-50), and were transferred after injection into 1/3x MR containing Gentamycin. Drop size was calibrated to about 7–8nL per injection. X. laevis allotatraploid and contains.L and.S allels for most genes.

Embryos injected with hormone-inducible constructs of (GFP-ΔN-tp63-GR and MCI-GR) (41, 52) were treated with 10µM Dexamethasone (Sigma-Aldrich/Merck #D4902) in ethanol from eight-cell stage until fixation. Ultrapure Ethanol (NeoFroxx #LC-8657.3) was used as vehicle control.

Morpholino oligonucleotides (MOs) were obtained from Gene Tools targeting *dmrt2*, *foxi1*, and *ubp1*. Foxi1 MO concentrations are indicated in the figures, dmrt2 and ubp1 MO concentrations are indicated in the list below.

mRNAs encoding, *foxi1 (25-100*ng/μl), *gfp-foxi1 (5-50*ng/μl*), ubp1 (50*ng/μl*) and dmrt2 (50*ng/μl) (this study; cloned using primers listed below into pCS107), *mcidas* (100ng/μl) (52), *foxj1* (100ng/μl) (62), *foxa1* (100ng/μl) (63, 64), *ΔN-tp63* (100ng/μl) (41) were injected together with *membrane-gfp* or *membrane-rfp* (at 50ng/μL) or *h2b-rfp* (at 30ng/μL) as lineage tracers. All mRNAs were prepared using the mMessage Machine kit using Sp6 (Invitrogen #AM1340) supplemented with RNAse Inhibitor (Promega #N251B).

The *foxi1::gfp-utrophin*, *foxi1ΔFoxi2BR::gfp-utrophin* and *a-tub::mscarletI* plasmids were purified using the Pure Yield midiprep kit (Promega #A2492) and injected at 5 ng/μl.

### Morpholino Sequences and doses

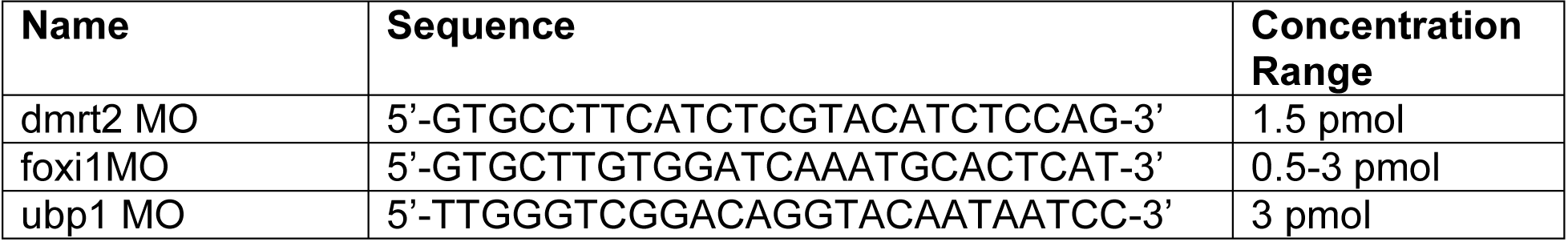

### Full length foxi1, gfp-foxi1, ubp1 and dmrt2 cloning (3’-5’)

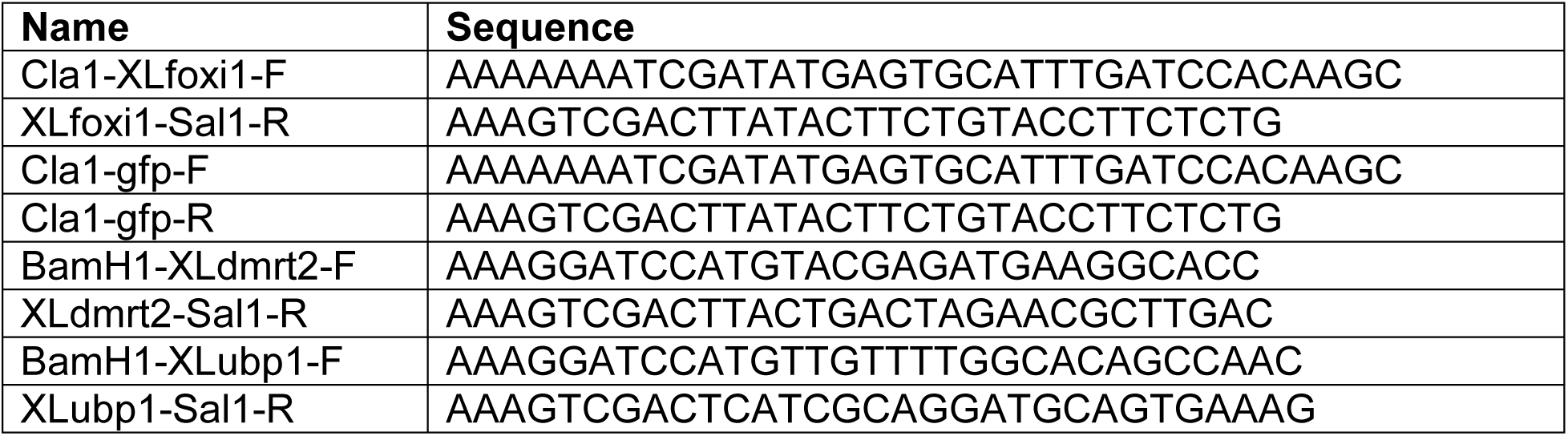

### *foxi1.S* reporter construct cloning and experiments

To generate the *foxi1.S::gfp-utrophin* reporter construct, genomic DNA was prepared from *X. laevis* using the phenol/chloroform DNA purification (ThermoFisher #15593031 and associated protocol). A 2.7 kb fragment (Fig.S1B,C) of the *foxi1.S* promoter was cloned using Easy-A Hi-Fi Cloning Enzyme (Agilent #600404) and primers listed in the table below. The PCR fragment was ligated using the pGEM-T Easy Vector System (Promega #A1360). The *foxi1.S* promoter sequence was subcloned into *a-tub::gfp-utrophin* (used in (22)) after removal of the *a-tub* promoter sequence using HiFi DNA Assembly (NEB #E2621S) and Q5 High-Fidelity DNA Polymerase (NEB #M0491S) kits. *foxi1ΔFoxi2BR::gfp-utrophin* reporter version (Fig. S1B,C) was generated using Q5 High-Fidelity DNA Polymerase and primers listed in the table below. The *a-tub::mscarletI* reporter was generated by replacing the *gfp-utrophin* sequence in *a-tub::gfp-utrophin* by the *mscarletI* sequence using HiFi DNA Assembly and Q5 High-Fidelity DNA Polymerase and primers listed below. Final construct sequences were analyzed by whole-plasmid nanopore sequencing.

### Cloning primers *foxi1.S* reporter (3’-5’)

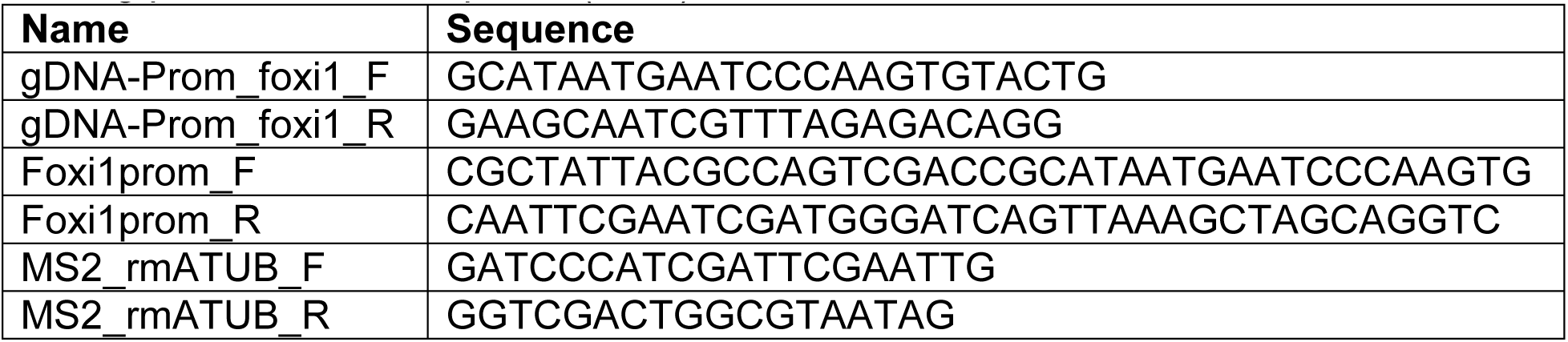
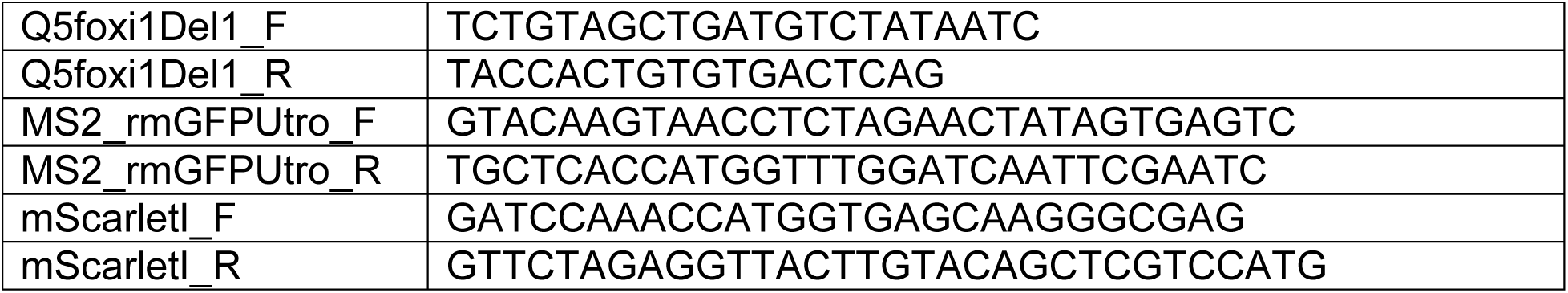

### Real-time quantitative PCR

Total RNA was extracted from 13 - 32 animal caps per sample at stage 10.5 - 11 using a standard Trizol (Invitrogen #15596026) protocol and used for cDNA synthesis with iScript cDNA Synthesis Kit (Bio-Rad #1708891). qPCR-reactions were conducted using Sso Advanced Universal SYBR Green Supermix (Bio-Rad #172-5275) on a CFX Connect Real-Time System (Bio-Rad) in 96-well PCR plates (Brand #781366). Expression values were normalized against two housekeeping control genes - EF1 and ODC (2^ΔΔpr^ method) (47). Results are presented as fold expression over average control sample values. Epidermal keratin (*krt14.4.S*; (65)) was used as pan-epidermal identity marker and *ubp1.L* was used as first selective ISC marker (3).

### qPCR primers (3’- 5’)

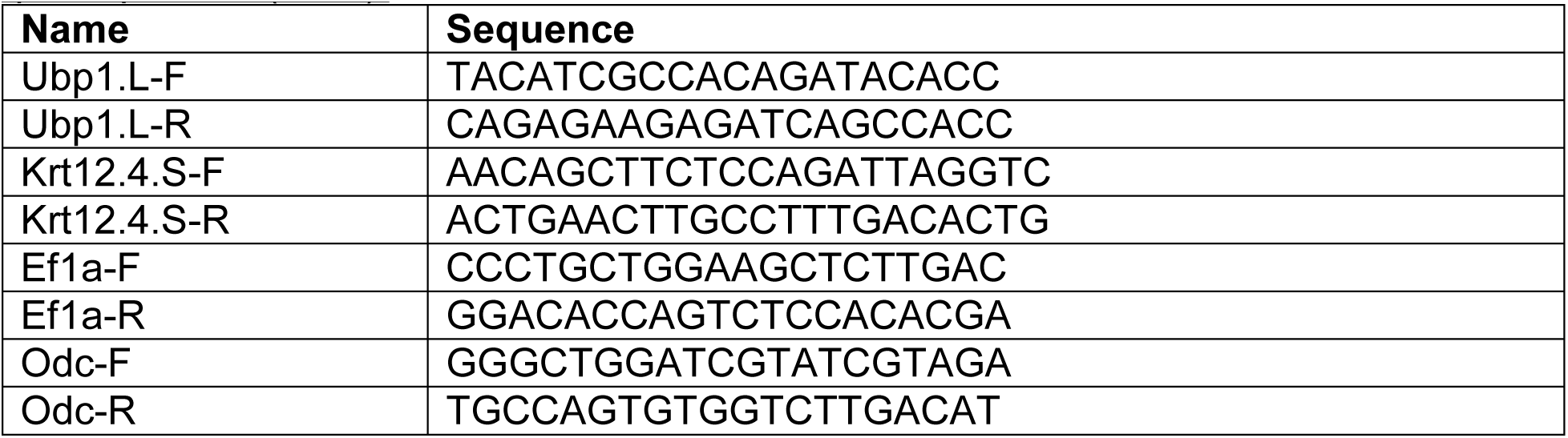

### Whole mount *in situ* hybridization and sections

For antisense *in situ* hybridization probes*, slc26a4*, *slc4a1*, *ubp1* and *dmrt2* fragments were cloned from whole-embryo cDNAs derived from stages between 3 and 30 using primers listed below (ISH-primers). All sequences were verified by Sanger sequencing. In addition, the following, previously published probes were used: *foxi1* (3), *foxj1* and *mcidas* (52, 62), *foxa1* (64), *tp63* (41), *atp6v1e1* (47) and *dll1* (22).

Embryos were fixed in MEMFA (100mM MOPS pH7.4, 2mM EGTA, 1mM MgSO4, 3.7% (v/v) Formaldehyde) overnight at 4°C and stored in 100% Ethanol at-20°C until used. DNAs were purified using the PureYield Midiprep kit (Promega #A2492) and were linearized before in vitro synthesis of anti-sense RNA probes using T7 or Sp6 polymerase (Promega, #P2077 and #P108G), RNAse inhibitor and dig-labeled rNTPs (Roche, #3359247910 and 11277057001). Embryos were in situ hybridized according to (66), bleached after staining with BM Purple (Roche #11442074001) and imaged. Sections were made after embedding in gelatin-albumin with Glutaraldehyde at 50-70μm as described in (67).

### Probe cloning primers (5’-3’)

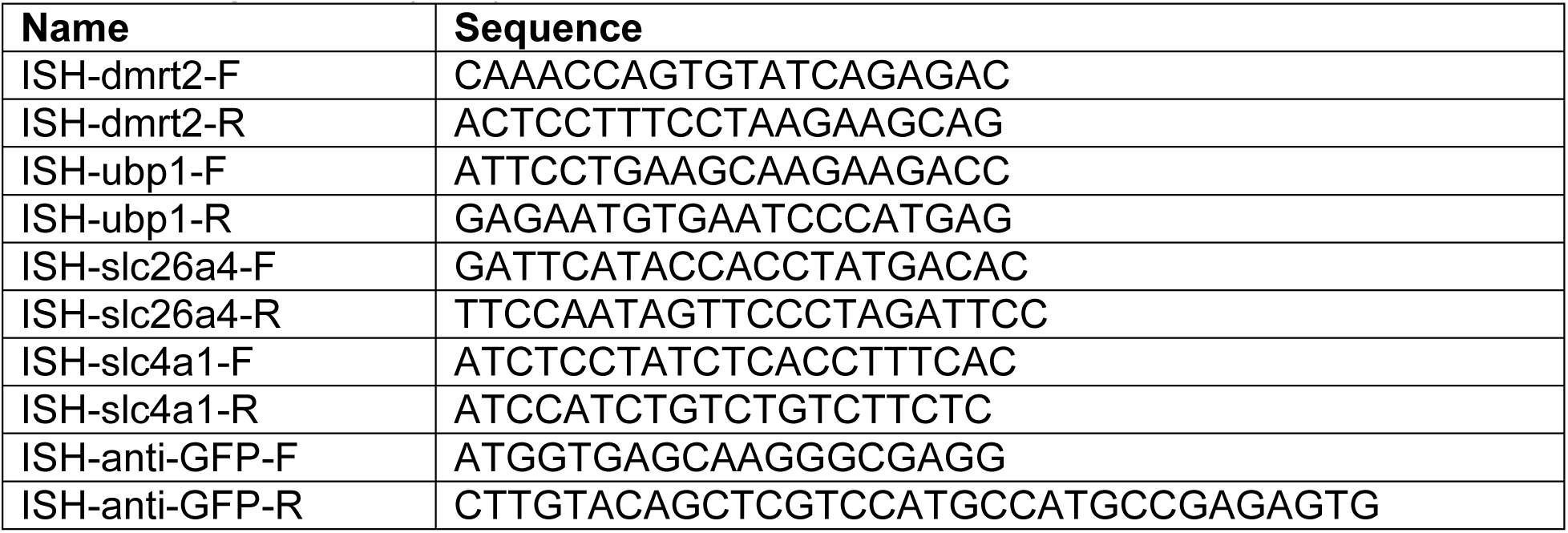

### TUNEL

Embryos were fixed at stage 9 - 10 in 1x MEMFA (100mM MOPS pH7.4, 2mM EGTA, 1mM MgSO4, 3.7% (v/v) Formaldehyde) overnight at 4°C or for 2h at RT, and stored in 100% Ethanol at - 20°C until use. Embryos were bleached before staining. TUNEL staining was performed as described in (22) using Terminal Deoxynucleotidyl Transferase Kit (Invitrogen #10533065), dig-UTP (Roche, #3359247910), anti-Digoxigenin AP Fab fragments (Roche, #11093274910) and NBT/BCIP (Roche, #11681451001). Staining was stopped with 100% Methanol (Roth, #8388.2), samples were then fixed briefly with 4% PFA (Roth, #0335.1) in PBS (Phosphate buffered saline, 10mM Na2HPO4, 1.8mM KH2PO4, 137mM NaCl, 2.7mM KCl) and imaged on a Zeiss Stemi508 with Axiocam208- color. Images were adjusted for color balance, brightness and contrast using Adobe Photoshop. Stage 43 tadpoles served as positive control samples for successful TUNEL staining.

### Evaluation of WMISH staining and morphological evaluations

Embryos were staged according to Nieuwkoop and Faber (1994) Normal Table of *Xenopus laevis* (Daudin). Garland Publishing Inc, New York ISBN 0-8153-1896-0. For the *foxi1* expression stage series wt embryos from multiple batches were mixed and at least 5 embryos per stage were assessed (Fig. 1D, S1A). Images of embryos after *in situ* hybridization and corresponding sections were imaged using a Zeiss AxioZoom setup, Zeiss AxioImager.Z1 or Zeiss Stemi508 with Axiocam208-color, and images were adjusted for color balance, brightness and contrast using Adobe Photoshop.

In Fig. 4C-G and S6D,E, embryos were quantified by selecting a region of interest (ROI) from each image of a stained embryo, and using the following FIJI/ImageJ adjustments processes to calculate the stained area: Color Deconvolution (H DAB), Convert to Mask, Analyze Particles. Macros used for this are available at www.github.com/sarahbowden/Imaging_Macros. In cases where the resulting files did not sufficiently represent the stained area of the ROI, this was generated manually with a combination of Brightness & Contrast, Color Deconvolution, Shadow, Threshold, or Color Thresholding techniques to select the most representative mask of stained regions. Quantified data was then plotted using a custom R script which performed the Wilcoxon

Rank Sum test to calculate p-values and generated plots using ggplot2. In Fig. 5A and S7D induction of expression was scored. In Fig. 5B *dll1* expression in the ventral epidermis was analyzed as normal or less (number of dots and expression intensity). In Fig. 5C, expression level differences observed between the uninjected control sides and manipulated sides of embryos were scored in whole mount embryos, while depicted sections are shown for clarity.

For analyses in Fig.3B and S4A, embryos injected with high dose of *foxi1* MO, cell morphology and cell size were evaluated for Fig. S4A (and delamination was confirmed in hemisected embryos) and skin lesions were evaluated for Fig. 3B.

### Immunofluorescence staining, *in situ* hybridization chain reaction and sample preparation

Whole *Xenopus* embryos, were fixed at indicated stages in 4% paraformaldehyde at 4°C overnight or 2h at room temperature, then washed 3x 15min with PBS, 2x 30min in PBST (0.1% Triton X-100 in PBS), and were blocked in PBST-CAS (90% PBS containing 0.1% Triton X-100, 10% CAS Blocking; ThermoFischer #00-8120) for 30min-1h at RT. A detailed protocol was described in (25).

Mouse anti-Acetylated-α-tubulin (Sigma/Merck #T6793) primary antibody (1:1000) was used to mark cilia / MCCs, Rabbit Anti-serotonin (Sigma/Merck #S5545) primary antibody (1:500) was used to mark SSCs, Rabbit Anti-GFP (Invitrogen #A11122) primary antibody (1:400) was used after HCR™ to boost *foxi1*::gfp-utrophin signal and applied at 4°C overnight. Secondary antibodies AlexaFluor-405-labeled goat anti-mouse (Invitrogen # A30104), AlexaFluor 405-labeled goat anti-rabbit antibody (Invitrogen #A31556) and AlexaFluor-488-labeled goat anti-rabbit antibody (Invitrogen #A11008) were used for 2 h at RT (1:250). Antibodies were applied in 100% CAS Blocking (ThermoFischer #00-8120). Actin was stained by incubation (30-120 min at room temperature) with AlexaFluor 405-labeled Phalloidin (1:800 in PBSt; Invitrogen #A30104), mucus-like compounds were stained by incubation (overnight at 4°C) with AlexaFluor-594-or-647-labeled or PNA (1:500-1000 in PBSt; Molecular Probes #L32459 and #L32460).

For HCR, *Xenopus* embryos were fixed at indicated stages in 10% MEMFA (100mM MOPS pH7.4, 2mM EGTA, 1mM MgSO4, 3.7% (v/v) Formaldehyde) for 1 h at RT, washed with PBSTw and stored in 100% Methanol at-20°C until use. HCR (hybridized chain reaction) was performed as previously described (68). *foxi1* probe and amplifiers were designed and obtained from Molecular Instruments, Inc. (https://www.molecularinstruments.com/). IF staining was performed on samples after HCR following the steps described above.

### Fluorescence imaging, image processing and analysis

Confocal imaging was conducted using either a Zeiss LSM880 or a Zeiss LSM980 microscope and Zeiss Zen software in the LIC and BiMiC imaging facilities. Confocal images were adjusted for channel brightness/contrast, Z-stack projections were generated and cell types were quantified based on their morphology using ImageJ (69).

For analyses in Fig. 3D and Fig. 5D, a detailed protocol for quantification of *Xenopus* epidermal cell type composition was published (25).

For analysis and comparison of fluorescent reporter construct activity on confocal micrographs (Fig. S5A) in ImageJ, z-projections were performed using the “sum-slices” function. Fluorescent intensities were color-coded using the function “lookup tables-> fire” (8 bit) in ImageJ. Induction of reporter expression in the endoderm (Fig. S5B) embryos were imaged using a Zeiss AxioImager.Z1 with Axiocam208-color camera. Induction was scored as positive when GFP fluorescence was detected in the vegetal half of the gastrula embryo. In some controls, activity was observed in involuting or animally positioned mesoderm, where maternal *foxi2* deposition occurs.

### Western blot analysis of GFP-Foxi1 levels

8-15 embryos per condition were collected and stored at-80 °C until use. Embryos were lysed in 100 µl of 1x Lysis Buffer (20 mM Tris-HCl pH8, 150 mM NaCl, 2 mM EDTA, 1x Protease Inhibitor Roche, #04693116001, 1% NP40 Sigma, #I8896) and smashed by pipetting up and down, then the samples were centrifuged at 4 °C at maximum speed for 15 minutes to remove yolk. 4x Laemmli Buffer (50 ml 4x buffer, 1 M Tris, pH 6.8, 4 g SDS, 20ml Glycerol, 10 ml 2-Mercaptoethanol, 0.1 g Bromophenol Blue) was added to the supernatant and the samples were cooked at 95 °C for 10 minutes.

10% separating gel (2.5 ml 4x Tris SDS (Roth, #2326), pH 8.8, 2 ml 40 % Acrylamide (Sigma, #A7802), 5.4 ml H_2_O, 40 µl TEMED (Roth, #2367.1), 100 µl 10 % APS (Roth, #9592.2)) was used, and a 4 % collecting gel (1.25 ml 4x Tris SDS, pH 6.8, 0.625 ml 40 % Acrylamide, 3.11 ml H_2_O, 50 µl TEMED, 50 µl 10 % APS)). 1x Running buffer (25 mM Tris-HCl, pH 8, 192 mM Glycine (Roth, #3187) in distilled water) was used for electrophoresis. 10 µl Precision Plus Protein Western C Standards Ladder (BioRad; #161-0376) and 20 µl of each sample were loaded for electrophoresis (120 V for 90 minutes).

Semi-dry transfer onto an activated PVDF membrane (Thermo Scientific, #88518) was conducted in 1x Towbin buffer with 0.1 % SDS for 60 minutes (25 mM Tris-base, 192 mM Glycine, 1 % SDS) using a PerfectBlue Semi-Dry Electroblotter Sedec M (VWR; 700-1220). Membranes and Gels were washed in 1x TBStw (100 mM Tris-base, 500 mM NaCl, 1 % Tween 20) at RT.

Membranes were blocked for at least 45 minutes using 5 % non-fat dry milk (Roth, #T145.3) in TBStw. The following primary antibodies were used at 1:1000 and incubated over night at 4 °C: Rabbit monoclonal anti-GFP (Abcam, #ab290) and, as a loading control, mouse monoclonal alpha-Tubulin (Thermo Scientific, DM1A #62204). The membrane was then washed in 1x TBStw for 4x 20 minutes. The following secondary antibodies were used at 1:5000 and incubated for 2 h at RT: HRP-linked Anit-Mouse IgG (Cell Signalling, #7076) and HRP-linked Anti-Rabbit IgG (Cell Signalling, #7074). The membrane was washed in 1x TBStw for 6x 10 minutes. Membranes were incubated with a mixture of 500 µl of Peroxide solution and 500 µl of Luminol/enhancer solution (both from Clarity™ Western ECL Substrate (BioRad, #170-5061) for 5 minutes in the dark at room temperature. Membranes were imaged using the Odyssey XF Imaging System by LI-COR. Afterwards, membranes were washed and stored in TBStw. Membranes were stripped for loading control (α-Tubulin) re-probing with stripping Buffer (2 % SDS, 50 mM Tris pH 6.8, 100 mM 2-Mercaptoethanol) for 1 h at 50 °C. The membranes were again rinsed in distilled water and washed 3x 5 minutes with TBStw. Brightness and contrast were adjusted in image J, and the ladder image was added to the chemi-luminescence membrane image.

### RNA-and ATAC-sequencing on *Xenopus* mucociliary organoids and bioinformatics analysis

Manipulations and bulk mRNA-seq used in this paper were generated and published here (41, 42) (GSE130448, n = 3 per time point and condition; GSE215373, n = 2 per time point and condition; GSE215419, n = 2 per time point; GSE262944, n = 2 per time point) or generated for this study (*foxi1* MO, *ubp1* MO, *dmrt2* MO on animal cap organoids at stages 20 and 32, n = 3 per time point and condition; GSE299718). scRNA-seq datasets were published here: (49, 53).

Data for Fig. 4B and S6A-C was generated from 10-15 pooled animal caps per sample and time point (3 replicates each), collected in Trizol for total RNA isolation. Library preparation and RNA-sequencing (150 base paired-end reads, min. 15 Mio. reads per sample) were performed in collaboration with the NIG, University Medical Center Göttingen using standard protocols described in (41, 42). RNA-seq data generated for this study was deposited at NCBI GEO under (GSE299718).

For Fig. 3A, data from (42) were used, TPM values from.L and.S allo-allels were added, and the resulting matrix was clustered using Z-values per line and galaxy.eu (ggplot2_heatmap2/3.1.3.1+galaxy). For Fig. S7A-C, log2-fold changes were calculated using galaxy.eu (DeSeq2/2.1.3+galaxy) and visualized using (ggplot2_heatmap2/3.1.3.1+galaxy). For Fig. S5C, the online tool associated with (49) was used (marionilab.cruk.cam.ac.uk/XenopusRegeneration). For Fig. S7E, the online tool associated with (53) was used (kleintools.hms.harvard.edu/tools/currentDatasetsList_xenopus_v2.html) to extract lineage-enriched transcripts and the heatmap was generated using galaxy.eu (ggplot2_heatmap2/3.1.3.1+galaxy).

For ATAC-seq sample generation, injected and control embryos were cultured until st. 8. Animal caps were dissected in 1x Modified Barth’s solution (MBS) and transferred to 0.5x MBS + Gentamycin (26). 2 organoids per condition and replicate were collected in PBS and ATAC-seq was performed as described in (70, 71). In short: Embryos were injected bilaterally in the animal hemisphere at the two-cell stage with 3pmol *foxi1* MO or remained uninjected, animal caps were prepared at st. 8, and organoids were collected upon the appearance of the dorsal lip in control embryos cultured in parallel to the organoids (st. 10). Organoids were transferred from MBS plates into a 1.5 mL low-bind microcentrifuge tube (Eppendorf #0030108051) containing 1 mL of ice-cold 1x PBS. Samples were spun at 500 g at 4 °C in the centrifuge for five minutes before removing the PBS and repeating the wash step with fresh ice-cold 1x PBS. 50 µL of ice-cold lysis buffer (10 mM Tris pH 7.4, 10 mM NaCl, 3 mM MgCl2, 0.1% (w/v) Igepal CA-630) and pipetted to break up samples. Samples were centrifuged at 500 g for 10 min at 4 °C and pellets were resuspended in 25 µL TD Buffer, 2.5 µL TDE1 Enzyme and 22.5 µL Nuclease-Free water (Illumina #20034198). Samples were pelleted to mix and incubated on a ThermoMixer at 37 °C, 700 rpm for 30 min. Following incubation, the samples were cleaned with MinElute Reaction Cleanup Kit (Qiagen, #28204), following manufacturer instructions and eluted into 11 µL Buffer EB.

Libraries were prepared in collaboration with the NIG, University Medical Center Göttingen. Quality was assessed with the Agilent Fragment Analyzer and prepared with the ATAC-seq Kit (Active Motif, #53150). Samples were sequenced in triplicate on an Illumina NovaSeq6000 with 150 nucleotide paired-end reads, totaling 50 million reads per sample.

Raw sequencing files were assessed for quality using FastQC (v0.11.9, Andrews, S. FastQC A Quality Control tool for High Throughput Sequence Data. http://www.bioinformatics.babraham.ac.uk/projects/fastqc/), and adapter sequences were removed with TrimGalore (v0.6.7, https://doi.org/10.5281/zenodo.7598955). Data were aligned to the *X. laevis* genome assembly v9.2 using BWA-MEM (v0.7.17, https://arxiv.org/abs/1303.3997). Mitochondrial reads were removed using Samtools (v1.21) (72), and peak calling was performed with the callpeak function of MACS2 (v2.2.7.1) (73). Differential analysis was performed with the bdgdiff function of MACS2 and Venn diagrams were generated with VennDiagram v1.7.3 in R v4.4.1. Heatmaps showing the average ATAC-seq signal were generated using deepTools (v3.5.4) (74). Peaks were annotated for the nearest *X. laevis* gene and transcription factor binding motifs with Homer (v4.11) (75), Plant-specific transcription factors were manually excluded from the lists of transcription factors. In Fig. 2E,F, each peak in the analysis was annotated to the nearest coding gene using Homer (v4.11). The peaks which were nearest to mucociliary-specific genes (specific for ionocytes, multiciliated cells, and basal cells published in (40, 41)) were then identified. Venn diagrams were then made for each mucociliary cell type, comparing the number of peaks lost, maintained, or gained for each cell type. Bioinformatic analyses were performed on the Galaxy / Europe platform (usegalaxy.eu) (76). ATAC-seq data generated for this study was deposited at NCBI GEO under (GSE280790).

### Quantification and statistical evaluation

Stacked bar graphs were generated in Microsoft Excel. Heatmaps and Venn diagrams were generated using the Galaxy Europe platform (usegalaxy.eu) and R. Statistical tests used and significance levels are indicated in the figure legends, and were conducted in Excel or R. Sample sizes for all experiments were chosen based on previous experience and used embryos derived from at least two different females. No randomization or blinding was applied.

### Use of shared controls

For some of the *in situ* and IF experiments shared controls were used in multiple graphs; all experiments are listed in the supplemental experiment log.

## References

1. Pou Casellas C, Pleguezuelos-Manzano C, Rookmaaker MB, Verhaar MC, Clevers H. Transcriptomic profile comparison reveals conservation of ionocytes across multiple organs. Sci Rep. 2023;13(1):3516.

2. Hulander M, Kiernan AE, Blomqvist SR, Carlsson P, Samuelsson EJ, Johansson BR, et al. Lack of pendrin expression leads to deafness and expansion of the endolymphatic compartment in inner ears of Foxi1 null mutant mice. Development. 2003;130(9):2013–25.

3. Quigley IK, Stubbs JL, Kintner C. Specification of ion transport cells in the Xenopus larval skin. Development. 2011;138(4):705–14.

4. Janicke M, Carney TJ, Hammerschmidt M. Foxi3 transcription factors and Notch signaling control the formation of skin ionocytes from epidermal precursors of the zebrafish embryo. Dev Biol. 2007;307(2):258–71.

5. Mir A, Kofron M, Zorn AM, Bajzer M, Haque M, Heasman J, et al. FoxI1e activates ectoderm formation and controls cell position in the Xenopus blastula. Development. 2007;134(4):779–88.

6. Cha SW, McAdams M, Kormish J, Wylie C, Kofron M. Foxi2 is an animally localized maternal mRNA in Xenopus, and an activator of the zygotic ectoderm activator Foxi1e. PLoS One. 2012;7(7):e41782.

7. Suri C, Haremaki T, Weinstein DC. Xema, a foxi-class gene expressed in the gastrula stage Xenopus ectoderm, is required for the suppression of mesendoderm. Development. 2005;132(12):2733–42.

8. Walentek P, Quigley IK. What we can learn from a tadpole about ciliopathies and airway diseases: Using systems biology in Xenopus to study cilia and mucociliary epithelia. Genesis. 2017;55(1-2).

9. Walentek P. Xenopus epidermal and endodermal epithelia as models for mucociliary epithelial evolution, disease, and metaplasia. Genesis. 2021;59(1-2):e23406.

10. Whitsett JA. Airway Epithelial Differentiation and Mucociliary Clearance. Ann Am Thorac Soc. 2018;15(Suppl 3):S143–s8.

11. Walentek P. Signaling Control of Mucociliary Epithelia: Stem Cells, Cell Fates, and the Plasticity of Cell Identity in Development and Disease. Cells Tissues Organs. 2022;211(6):736–53.

12. Luan X, Henao Romero N, Campanucci VA, Le Y, Mustofa J, Tam JS, et al. Pulmonary Ionocytes Regulate Airway Surface Liquid pH in Primary Human Bronchial Epithelial Cells. Am J Respir Crit Care Med. 2024.

13. Montoro DT, Haber AL, Biton M, Vinarsky V, Lin B, Birket SE, et al. A revised airway epithelial hierarchy includes CFTR-expressing ionocytes. Nature. 2018;560(7718):319-24.

14. Blomqvist SR, Vidarsson H, Fitzgerald S, Johansson BR, Ollerstam A, Brown R, et al. Distal renal tubular acidosis in mice that lack the forkhead transcription factor Foxi1. J Clin Invest. 2004;113(11):1560–70.

15. Blomqvist SR, Vidarsson H, Soder O, Enerback S. Epididymal expression of the forkhead transcription factor Foxi1 is required for male fertility. EMBO J. 2006;25(17):4131–41.

16. Yang T, Vidarsson H, Rodrigo-Blomqvist S, Rosengren SS, Enerback S, Smith RJ. Transcriptional control of SLC26A4 is involved in Pendred syndrome and nonsyndromic enlargement of vestibular aqueduct (DFNB4). Am J Hum Genet. 2007;80(6):1055–63.

17. Lindgren D, Eriksson P, Krawczyk K, Nilsson H, Hansson J, Veerla S, et al. Cell-Type-Specific Gene Programs of the Normal Human Nephron Define Kidney Cancer Subtypes. Cell Rep. 2017;20(6):1476–89.

18. Simbolo M, Centonze G, Gkountakos A, Monti V, Maisonneuve P, Golovco S, et al. Characterization of two transcriptomic subtypes of marker-null large cell carcinoma of the lung suggests different origin and potential new therapeutic perspectives. Virchows Arch. 2024;484(5):777–88.

19. Skala SL, Wang X, Zhang Y, Mannan R, Wang L, Narayanan SP, et al. Next-generation RNA Sequencing-based Biomarker Characterization of Chromophobe Renal Cell Carcinoma and Related Oncocytic Neoplasms. Eur Urol. 2020;78(1):63–74.

20. Yamada Y, Belharazem-Vitacolonnna D, Bohnenberger H, Weiss C, Matsui N, Kriegsmann M, et al. Pulmonary cancers across different histotypes share hybrid tuft cell/ionocyte-like molecular features and potentially druggable vulnerabilities. Cell Death Dis. 2022;13(11):979.

21. Hendrickson CL, Blitz IL, Hussein A, Paraiso KD, Cho J, Klymkowsky MW, et al. Foxi2 and Sox3 are master regulators controlling ectoderm germ layer specification. bioRxiv. 2025.

22. Tasca A, Helmstadter M, Brislinger MM, Haas M, Mitchell B, Walentek P. Notch signaling induces either apoptosis or cell fate change in multiciliated cells during mucociliary tissue remodeling. Dev Cell. 2021;56(4):525–39 e6.

23. Ventrella R, Kim SK, Sheridan J, Grata A, Bresteau E, Hassan OA, et al. Bidirectional multiciliated cell extrusion is controlled by Notch-driven basal extrusion and Piezo1-driven apical extrusion. Development. 2023;150(17).

24. Yan J, Xu L, Crawford G, Wang Z, Burgess SM. The forkhead transcription factor FoxI1 remains bound to condensed mitotic chromosomes and stably remodels chromatin structure. Mol Cell Biol. 2006;26(1):155–68.

25. Walentek P. Manipulating and Analyzing Cell Type Composition of the Xenopus Mucociliary Epidermis. Methods Mol Biol. 2018;1865:251–63.

26. Sive HL, Grainger RM, Harland RM. Animal Cap Isolation from Xenopus laevis. CSH Protoc. 2007;2007:pdb.prot4744.

27. Luo T, Zhang Y, Khadka D, Rangarajan J, Cho KW, Sargent TD. Regulatory targets for transcription factor AP2 in Xenopus embryos. Dev Growth Differ. 2005;47(6):403–13.

28. Zhang Y, Luo T, Sargent TD. Expression of TFAP2beta and TFAP2gamma genes in Xenopus laevis. Gene Expr Patterns. 2006;6(6):589–95.

29. Ma L, Hocking JC, Hehr CL, Schuurmans C, McFarlane S. Zac1 promotes a Müller glial cell fate and interferes with retinal ganglion cell differentiation in Xenopus retina. Dev Dyn. 2007;236(1):192–202.

30. Schweickert A, Steinbeisser H, Blum M. Differential gene expression of Xenopus Pitx1, Pitx2b and Pitx2c during cement gland, stomodeum and pituitary development. Mech Dev. 2001;107(1-2):191–4.

31. Giudetti G, Giannaccini M, Biasci D, Mariotti S, Degl’innocenti A, Perrotta M, et al. Characterization of the Rx1-dependent transcriptome during early retinal development. Dev Dyn. 2014;243(10):1352–61.

32. Ray H, Chang C. The transcription factor Hypermethylated in Cancer 1 (Hic1) regulates neural crest migration via interaction with Wnt signaling. Dev Biol. 2020;463(2):169–81.

33. Afouda BA, Ciau-Uitz A, Patient R. GATA4, 5 and 6 mediate TGFbeta maintenance of endodermal gene expression in Xenopus embryos. Development. 2005;132(4):763–74.

34. Smith JC, Price BM, Green JB, Weigel D, Herrmann BG. Expression of a Xenopus homolog of Brachyury (T) is an immediate-early response to mesoderm induction. Cell. 1991;67(1):79–87.

35. Hopwood ND, Pluck A, Gurdon JB. MyoD expression in the forming somites is an early response to mesoderm induction in Xenopus embryos. Embo j. 1989;8(11):3409–17.

36. Jerabek S, Merino F, Schöler HR, Cojocaru V. OCT4: dynamic DNA binding pioneers stem cell pluripotency. Biochim Biophys Acta. 2014;1839(3):138–54.

37. Mistri TK, Devasia AG, Chu LT, Ng WP, Halbritter F, Colby D, et al. Selective influence of Sox2 on POU transcription factor binding in embryonic and neural stem cells. EMBO Rep. 2015;16(9):1177–91.

38. Wills AE, Choi VM, Bennett MJ, Khokha MK, Harland RM. BMP antagonists and FGF signaling contribute to different domains of the neural plate in Xenopus. Dev Biol. 2010;337(2):335–50.

39. Deblandre GA, Wettstein DA, Koyano-Nakagawa N, Kintner C. A two-step mechanism generates the spacing pattern of the ciliated cells in the skin of Xenopus embryos. Development. 1999;126(21):4715–28.

40. Quigley IK, Kintner C. Rfx2 Stabilizes Foxj1 Binding at Chromatin Loops to Enable Multiciliated Cell Gene Expression. PLoS Genet. 2017;13(1):e1006538.

41. Haas M, Gomez Vazquez JL, Sun DI, Tran HT, Brislinger M, Tasca A, et al. DeltaN-Tp63 Mediates Wnt/beta-Catenin-Induced Inhibition of Differentiation in Basal Stem Cells of Mucociliary Epithelia. Cell Rep. 2019;28(13):3338–52 e6.

42. Brislinger-Engelhardt MM, Lorenz F, Haas M, Bowden S, Tasca A, Kreutz C, et al. Temporal Notch signaling regulates mucociliary cell fates through Hes-mediated competitive de-repression. bioRxiv. 2023.

43. Dubaissi E, Papalopulu N. Embryonic frog epidermis: a model for the study of cell-cell interactions in the development of mucociliary disease. Dis Model Mech. 2011;4(2):179–92.

44. Werth M, Schmidt-Ott KM, Leete T, Qiu A, Hinze C, Viltard M, et al. Transcription factor TFCP2L1 patterns cells in the mouse kidney collecting ducts. Elife. 2017;6.

45. Kurth I, Hentschke M, Hentschke S, Borgmeyer U, Gal A, Hübner CA. The forkhead transcription factor Foxi1 directly activates the AE4 promoter. Biochem J. 2006;393(Pt 1):277–83.

46. Valencia JE, Peter IS. Combinatorial regulatory states define cell fate diversity during embryogenesis. Nature Communications. 2024;15(1):6841.

47. Walentek P, Beyer T, Hagenlocher C, Muller C, Feistel K, Schweickert A, et al. ATP4a is required for development and function of the Xenopus mucociliary epidermis - a potential model to study proton pump inhibitor-associated pneumonia. Dev Biol. 2015;408(2):292–304.

48. Wu ST, Feng Y, Song R, Qi Y, Li L, Lu D, et al. Foxp1 Is Required for Renal Intercalated Cell Differentiation and Acid-Base Regulation. J Am Soc Nephrol. 2024;35(5):533–48.

49. Aztekin C, Hiscock TW, Marioni JC, Gurdon JB, Simons BD, Jullien J. Identification of a regeneration-organizing cell in the Xenopus tail. Science. 2019;364(6441):653-8.

50. Mukherjee M, DeRiso J, Janga M, Fogarty E, Surendran K. Foxi1 inactivation rescues loss of principal cell fate selection in Hes1-deficient kidneys but does not ensure maintenance of principal cell gene expression. Dev Biol. 2020;466(1-2):1–11.

51. Cibois M, Luxardi G, Chevalier B, Thome V, Mercey O, Zaragosi LE, et al. BMP signalling controls the construction of vertebrate mucociliary epithelia. Development. 2015;142(13):2352–63.

52. Stubbs JL, Vladar EK, Axelrod JD, Kintner C. Multicilin promotes centriole assembly and ciliogenesis during multiciliate cell differentiation. Nat Cell Biol. 2012;14(2):140–7.

53. Briggs JA, Weinreb C, Wagner DE, Megason S, Peshkin L, Kirschner MW, et al. The dynamics of gene expression in vertebrate embryogenesis at single-cell resolution. Science. 2018;360(6392).

54. Edri T, Cohen D, Shabtai Y, Fainsod A. Alcohol induces neural tube defects by reducing retinoic acid signaling and promoting neural plate expansion. Front Cell Dev Biol. 2023;11:1282273.

55. Hoffman TL, Javier AL, Campeau SA, Knight RD, Schilling TF. Tfap2 transcription factors in zebrafish neural crest development and ectodermal evolution. J Exp Zool B Mol Dev Evol. 2007;308(5):679–91.

56. Tremblay M, Sanchez-Ferras O, Bouchard M. GATA transcription factors in development and disease. Development. 2018;145(20).

57. Hou L, Srivastava Y, Jauch R. Molecular basis for the genome engagement by Sox proteins. Semin Cell Dev Biol. 2017;63:2–12.

58. Walentek P. Mucociliary cell type compositions - bridging the gap between genes and emergent tissue functions. Cells Dev. 2025:204019.

59. Fisher M, James-Zorn C, Ponferrada V, Bell AJ, Sundararaj N, Segerdell E, et al. Xenbase: key features and resources of the Xenopus model organism knowledgebase. Genetics. 2023;224(1).

60. Sive HL, Grainger RM, Harland RM. Xenopus laevis In Vitro Fertilization and Natural Mating Methods. CSH Protoc. 2007;2007:pdb.prot4737.

61. Sive HL, Grainger RM, Harland RM. Microinjection of Xenopus embryos. Cold Spring Harb Protoc. 2010;2010(12):pdb.ip81.

62. Stubbs JL, Oishi I, Izpisúa Belmonte JC, Kintner C. The forkhead protein Foxj1 specifies node-like cilia in Xenopus and zebrafish embryos. Nat Genet. 2008;40(12):1454–60.

63. Dubaissi E, Rousseau K, Lea R, Soto X, Nardeosingh S, Schweickert A, et al. A secretory cell type develops alongside multiciliated cells, ionocytes and goblet cells, and provides a protective, anti-infective function in the frog embryonic mucociliary epidermis. Development. 2014;141(7):1514–25.

64. Walentek P, Bogusch S, Thumberger T, Vick P, Dubaissi E, Beyer T, et al. A novel serotonin-secreting cell type regulates ciliary motility in the mucociliary epidermis of Xenopus tadpoles. Development. 2014;141(7):1526–33.

65. Vetrova AA, Kupaeva DM, Kizenko A, Lebedeva TS, Walentek P, Tsikolia N, et al. The evolutionary history of Brachyury genes in Hydrozoa involves duplications, divergence, and neofunctionalization. Scientific Reports. 2023;13(1):9382.

66. Harland RM. In situ hybridization: an improved whole-mount method for Xenopus embryos. Methods Cell Biol. 1991;36:685–95.

67. Walentek P, Beyer T, Thumberger T, Schweickert A, Blum M. ATP4a is required for Wnt-dependent Foxj1 expression and leftward flow in Xenopus left-right development. Cell Rep. 2012;1(5):516–27.

68. Huber PB, LaBonne C. Small molecule-mediated reprogramming of Xenopus blastula stem cells to a neural crest state. Developmental Biology. 2024;505:34–41.

69. Schindelin J, Arganda-Carreras I, Frise E, Kaynig V, Longair M, Pietzsch T, et al. Fiji: an open-source platform for biological-image analysis. Nat Methods. 2012;9(7):676–82.

70. Esmaeili M, Blythe SA, Tobias JW, Zhang K, Yang J, Klein PS. Chromatin accessibility and histone acetylation in the regulation of competence in early development. Dev Biol. 2020;462(1):20–35.

71. Buenrostro JD, Giresi PG, Zaba LC, Chang HY, Greenleaf WJ. Transposition of native chromatin for fast and sensitive epigenomic profiling of open chromatin, DNA-binding proteins and nucleosome position. Nat Methods. 2013;10(12):1213–8.

72. Danecek P, Bonfield JK, Liddle J, Marshall J, Ohan V, Pollard MO, et al. Twelve years of SAMtools and BCFtools. GigaScience. 2021;10(2).

73. Feng J, Liu T, Qin B, Zhang Y, Liu XS. Identifying ChIP-seq enrichment using MACS. Nat Protoc. 2012;7(9):1728–40.

74. Ramírez F, Ryan DP, Grüning B, Bhardwaj V, Kilpert F, Richter AS, et al. deepTools2: a next generation web server for deep-sequencing data analysis. Nucleic Acids Research. 2016;44(W1):W160–W5.

75. Heinz S, Benner C, Spann N, Bertolino E, Lin YC, Laslo P, et al. Simple combinations of lineage-determining transcription factors prime cis-regulatory elements required for macrophage and B cell identities. Mol Cell. 2010;38(4):576–89.

76. Community TG. The Galaxy platform for accessible, reproducible, and collaborative data analyses: 2024 update. Nucleic Acids Research. 2024;52(W1):W83–W94.

