## Supplemental Figures 1-7 for "Foxi1 regulates multipotent mucociliary progenitors and ionocyte specification through transcriptional and epigenetic mechanisms"

**Supplemental figures and legends:**

**Bowden, Brislinger, Hansen et al. Figure S1**

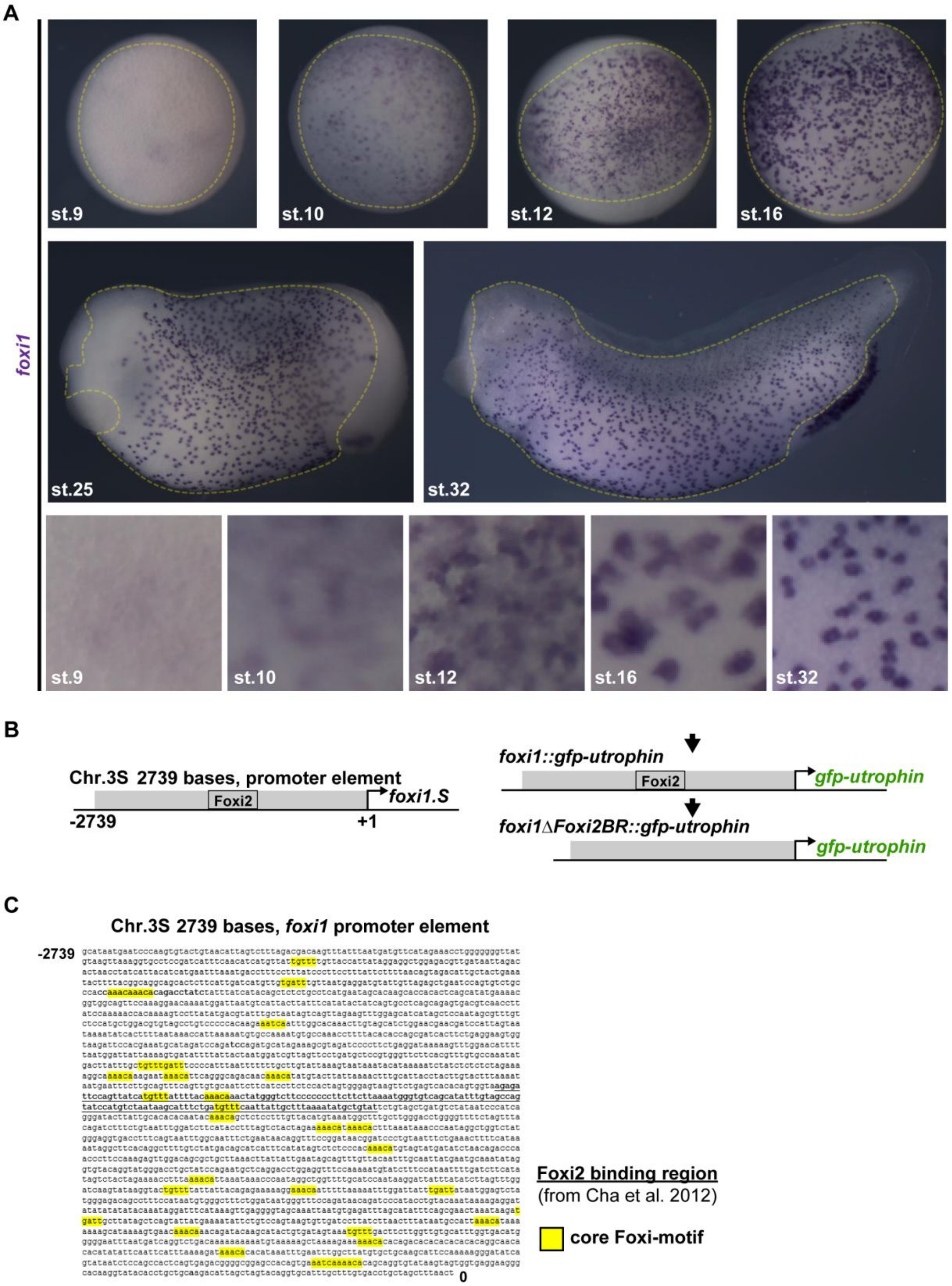

**Figure S1: Early Foxi1 expression and reporter constructs**

**(A)** WMISH expression analysis of *foxi1* across mucociliary epidermis development stages (st. 9 - 32). St. 9, 10 = animal views; st. 12, 16 = ventral views; st. 25, 32 = lateral views, anterior to the left. Ectodermal (st. 9 – 10) and epidermal (st. 12 – 32) are outlined in yellow. Epidermal  Bottom row panels = magnified views of epidermal areas. **(B,C)** Generation and promoter sequences of *foxi1::gfp-utrophin* or *foxi1ΔFoxi2BR::gfp-utrophin* reporters. **(B)** Schematic representation of cloned genomic *foxi1.S* promoter locus (grey box) and position of Foxi2 binding region determined in Cha et al., 2012 (black outlined box). **(C)** Promoter sequence with indicated predicted core Foxi binding-motifs (yellow) and Foxi2 binding region (bold, underscored).

Bowden, Brislinger, Hansen et al. Figure S2

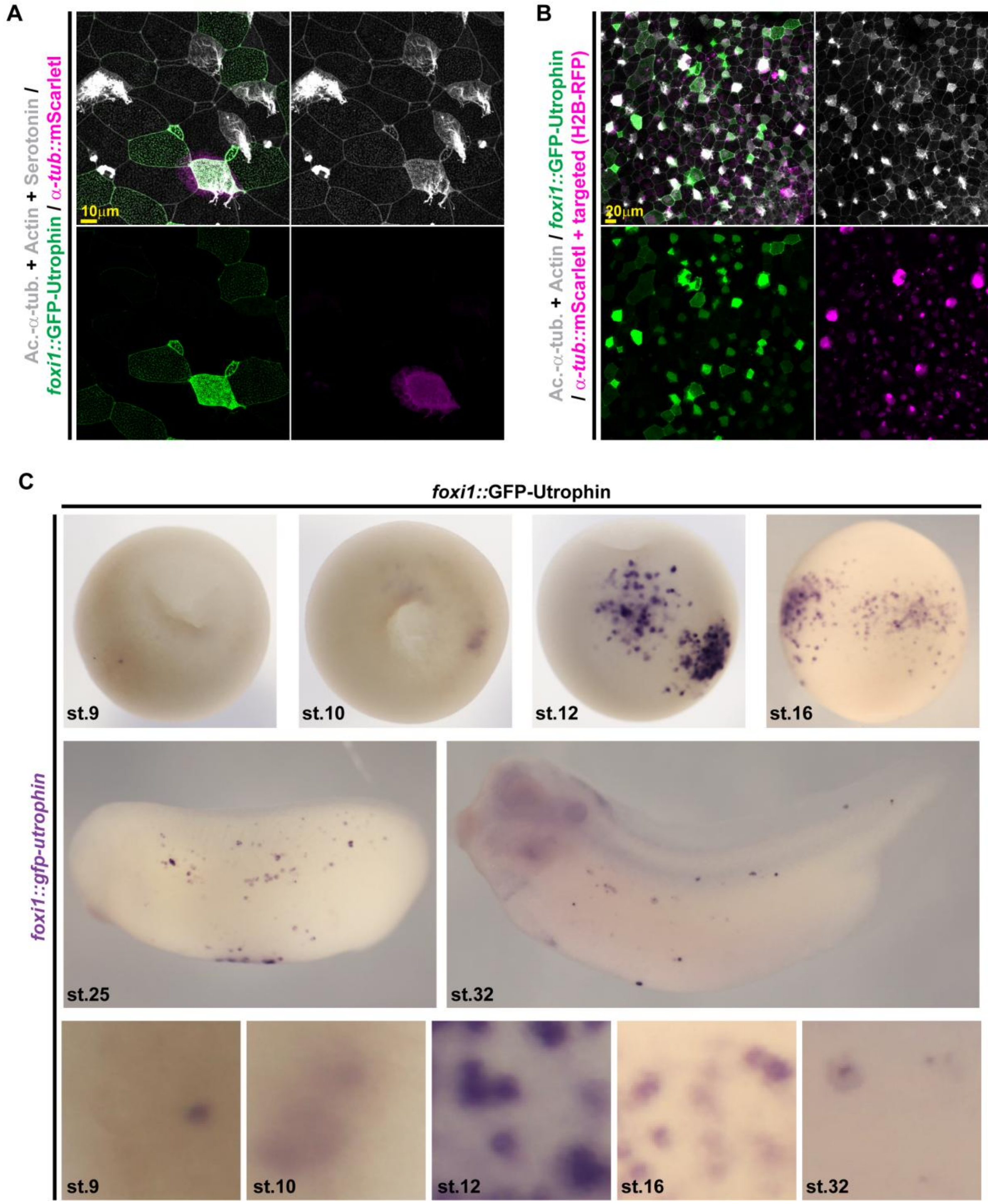

**Figure S2: Characterization of the foxi1 reporter**

**(A,B)** IF of embryos injected with *foxi1::gfp-utrophin* (green) (n = 12 embryos) and $\alpha$-tub.::mscarletI (magenta) (n = 9 embryos) reporters at st. 32 stained for Acetylated-$\alpha$-tubulin (Ac.-$\alpha$-tub., cilia, grey), F-actin (Actin, cell borders and morphology, grey), and serotonin (SSCs, grey) in **(A)**; or for Acetylated-$\alpha$-tubulin (Ac.-$\alpha$-tub., cilia, grey) and F-actin (Actin, cell borders and morphology, grey), in **(B)**. In **(B)** targeted cells were identified by nuclear RFP expression (H2B-RFP, magenta). **(C)** WMISH expression analysis of *foxi1::gfp-utrophin* (stained for *gfp* transcripts) across mucociliary epidermis development stages (st. 9 - 32). St. 9, 10 = animal views; st. 12, 16 = ventral views; st. 25, 32 = lateral views. Bottom row panels = magnified views of epidermal areas. Related to sections shown in Fig. 1D. st. 9 n = 17; st. 10 n = 19; st. 12 n = 16 ; st. 16 n = 14 ; st. 25 n = 14 ; st. 32 n = 19 embryos.

Bowden, Brislinger, Hansen et al. Figure S3

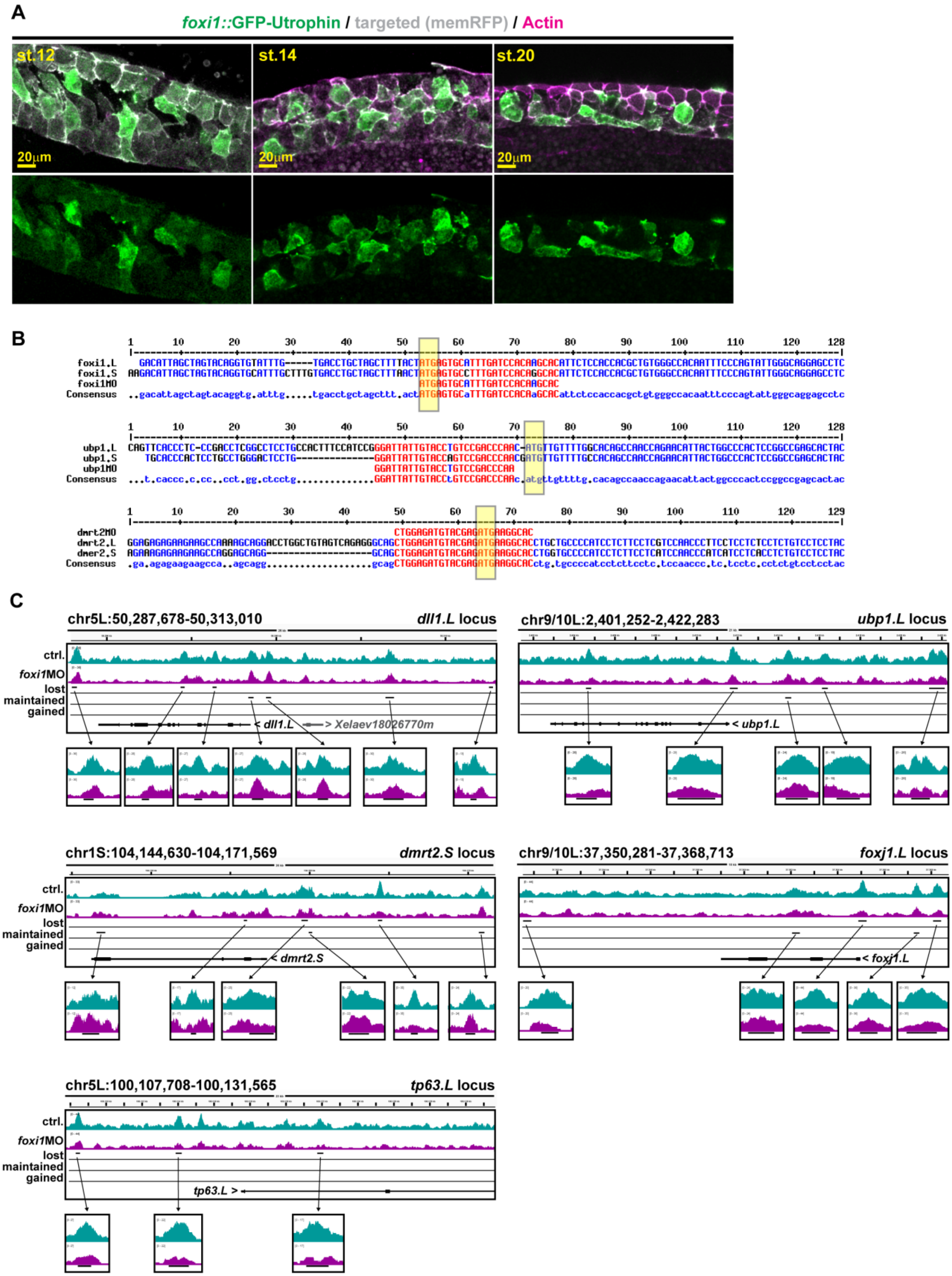

**Figure S3: Foxi1 reporter expression, morpholino targets, ATAC-profiles**

**(A)** IF for *foxi1::gfp-utrophin* reporter (green) and F-actin (Actin, cell borders and morphology, magenta) at st. 12 - 20 on hemisected embryos. Targeted cells were identified by membrane RFP expression (mRFP, grey). Related to sections shown in Fig. 1E. st. 12 n = 5 ; st. 14 n = 4 ; st. 20 n = 5 embryos. **(B)** Alignment of MO-target sequences in foxi1, ubp1 and dmrt2 transcripts. ATG start-codons are indicated by yellow boxes. Generated with http://multalin.toulouse.inra.fr. **(C)** Distribution of accessible regions around genes required for development and cell fates specification in the embryonic mucociliary epidermis of *Xenopus*. Lost, maintained and gained tracks as generated by MACS2 bdgdiff analysis and visualized in IGV: *dll1.L; ubp1.L; dmrt2.S; foxj1.L;* and *tp63.L.* Turquoise track = control (ctrl.) and purple track = morphant (*foxi1* MO). n = 2 organoids per condition and replicate. 3 replicates. Related to Fig. 2D.

Bowden, Brislinger,Hansen et al. Figure S4

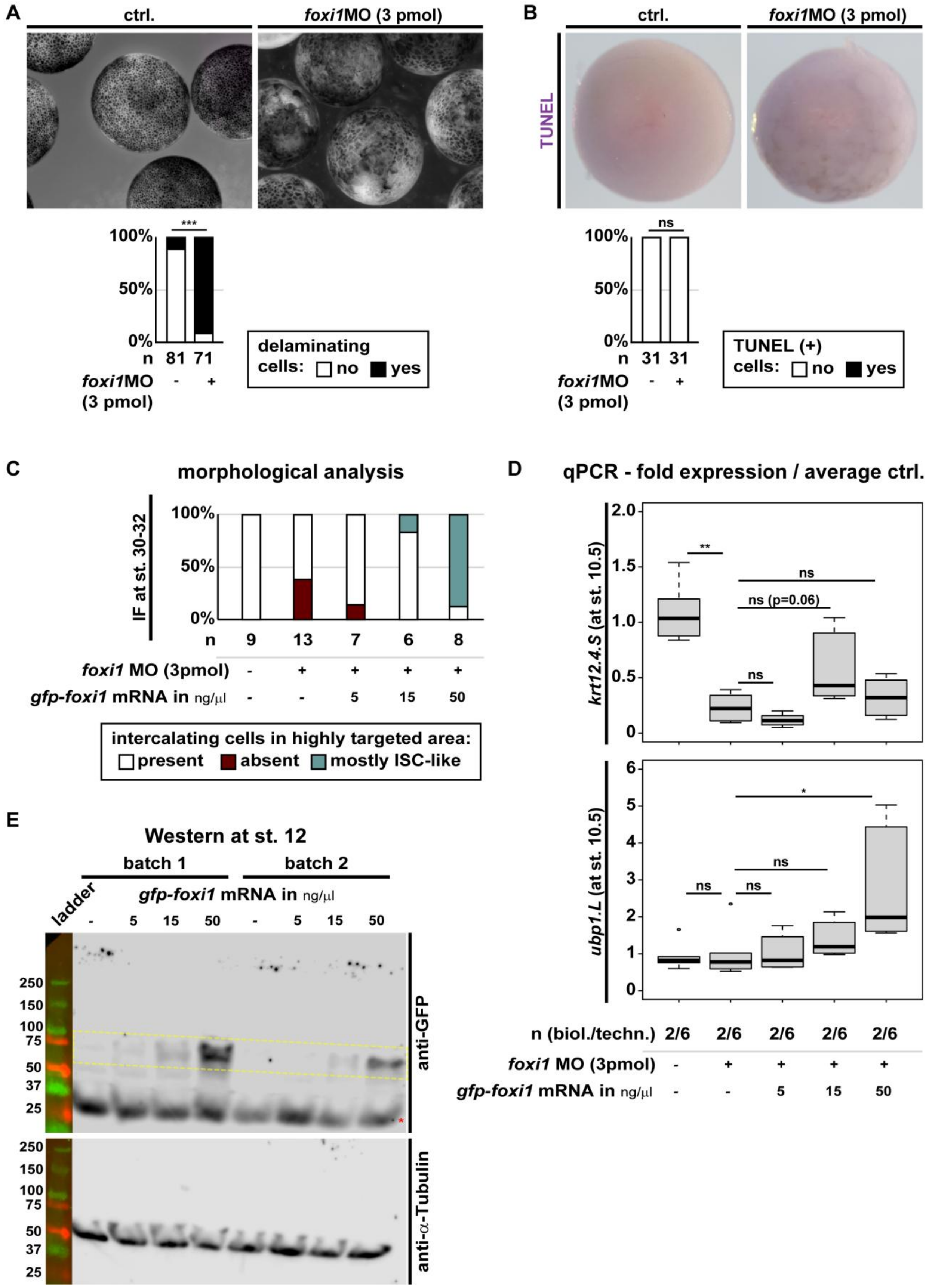

**Figure S4: Concentration-dependent effects of Foxi1 manipulations**

**(A)** Representative brightfield images of controls (ctrl.) and embryos (animal views) after *foxi1* MO (3 pmol) injection at st. 8. Morphants showed enlarged cells and delamination of animal cells into the blastocoel. Quantification of results shown in the graph. Delamination events were scored based on morphological analysis. n = number of embryos. Chi² test: $p < 0.001$ = ***. **(B)** TUNEL staining to identify apoptotic cells. Representative images of controls (ctrl.) and embryos (animal views) after *foxi1* MO (3 pmol) injection at st. 9-10. Quantification of results shown in the graph. n = number of embryos. Chi² test: $p > 0.05$ = ns. **(C)** Quantification of results depicted in Fig. 3C. Samples were analyzed for presence (white) or absence (dark red) if intercalating cells in highly targeted areas as well as for the presence of ISC-like cells (turquois). **(D)** qPCR on pooled uninjected control organoids and after *foxi1* MO (3 pmol) with or without co-injected *gfp-foxi1* at 5, 15 or 50 ng/μl. The epidermal competence gene *krt12.4.S* and the definitive ISC marker *ubp1* show differential dose-dependent reactions to *foxi1* manipulations. Student T-test: $p > 0.05$ = ns; $p < 0.05$ = *; $p < 0.01$ = **. n = number of biological and technical replicates. **(E)** Western blot analysis of GFP-Foxi1 overexpression (anti-GFP) levels in lysates from pooled whole embryos at stage 12 in uninjected controls and embryos injected with *gfp-foxi1* at 5, 15 or 50 ng/μl. Two different batches (biological replicates) are shown. Predicted size of GFP-Foxi1 ca. 68 kDa, specific bands are indicated by yellow box, unspecific band indicated by red asterisk. Anti-Tubulin is used as loading control.

**Bowden, Brislinger, Hansen et al. Figure S5**

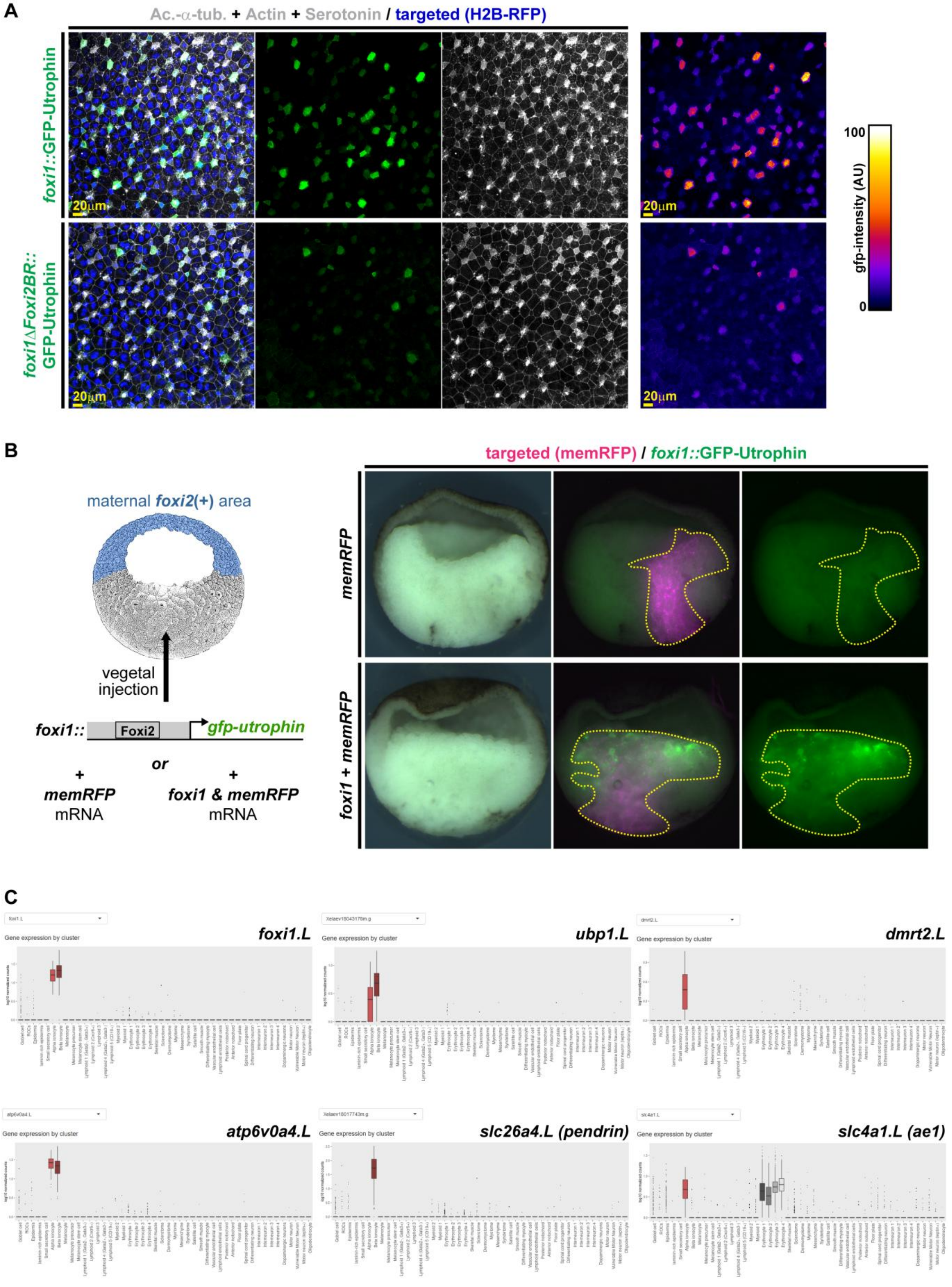

**Figure S5: Foxi1 regulates its own expression and Dmrt2 is expressed only in α-ISCs**

**(A)** Semi-quantitative IF analysis of embryos injected with *foxi1::gfp-utrophin* (n = 12 embryos) or *foxi1ΔFoxi2BR::gfp-utrophin* (n = 12 embryos) reporters (green) at st. 32 stained for Acetylated-α-tubulin (Ac.-α-tub., cilia, grey), F-actin (Actin, cell borders and morphology, grey), and serotonin (SSCs, grey) at st. 32. Targeted cells were identified by nuclear RFP expression (H2B-RFP, blue). Right panels show false-color of GFP fluorescence intensity. **(B)** Brightfield and epifluorescence images of hemisected st. 11 gastrula embryos injected vegetally with *foxi1::gfp-utrophin* (green), membrane RFP (memRFP; magenta) as control (*memRFP*) or with additional co-injection of *foxi1* mRNA (*foxi1* + *memRFP*). Right panels show false-color of GFP fluorescence intensity. Induction was scored as positive when GFP was detected in areas below the equator (mesendoderm). Ctrl. n = 7 induced, 26 non-induced; *foxi1* mRNA = 26 induced, 11 non-induced. Embryos are shown dorsal to the left and animal up. **(C)** Boxplots of ISC gene expression from scRNA-seq data published in Aztekin et al., 2019. Visualization was generated using the published online tool: marionilab.cruk.cam.ac.uk/XenopusRegeneration.

Bowden, Brislinger, Hansen et al. Figure S6

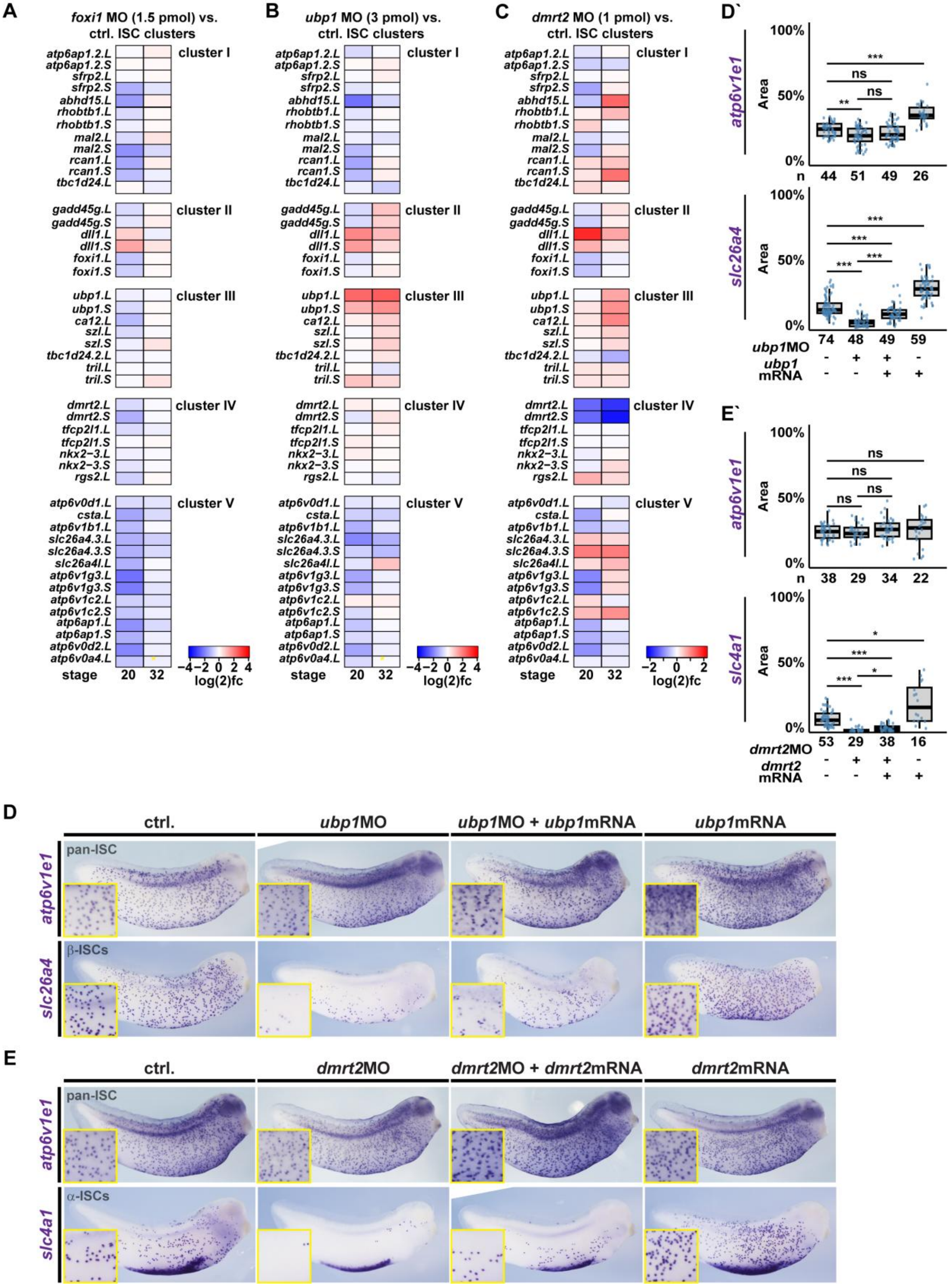

**Figure S6: Foxi1, Ubp1 and Dmrt2 differentially regulate ISC specification**
**(A,B,C)** Effects of Foxi1 (*foxi1* MO, 1.5 pmol; **A**), Ubp1 (*ubp1* MO, 3 pmol; **B**) or Dmrt2 (*dmrt2* MO, 1 pmol; **C**) knockdown on core ISC gene expression stages 20 and 32. RNA-seq on mucociliary organoids. Heatmaps depict log2-fold change values derived from DEseq2. **(D,E)** Analysis of effects by WMISH at st. 29 - 32 against *atp6v1e1* and *foxi1* (pan-ISC markers), *ubp1* and *slc25a4/pendrin* (β-ISC markets), and *dmrt2* and *slc4a1/ae1* (α-ISC markers) after Ubp1 (*ubp1* MO, 3 pmol) or Dmrt2 (*dmrt2* MO, 1 pmol) knockdown, rescue and overexpression (by mRNA injections: 50 ng/ul *ubp1*; 25-50 ng/ul *dmrt2*). Representative images and quantification of results are depicted. n = number of embryos analyzed per condition. Wilcoxon Rank Sum test: $p > 0.05$ = ns; $p < 0.05$ = *; $p < 0.01$ = **; $p < 0.001$ = ***.

**Bowden, Brislinger, Hansen et al. Figure S7**

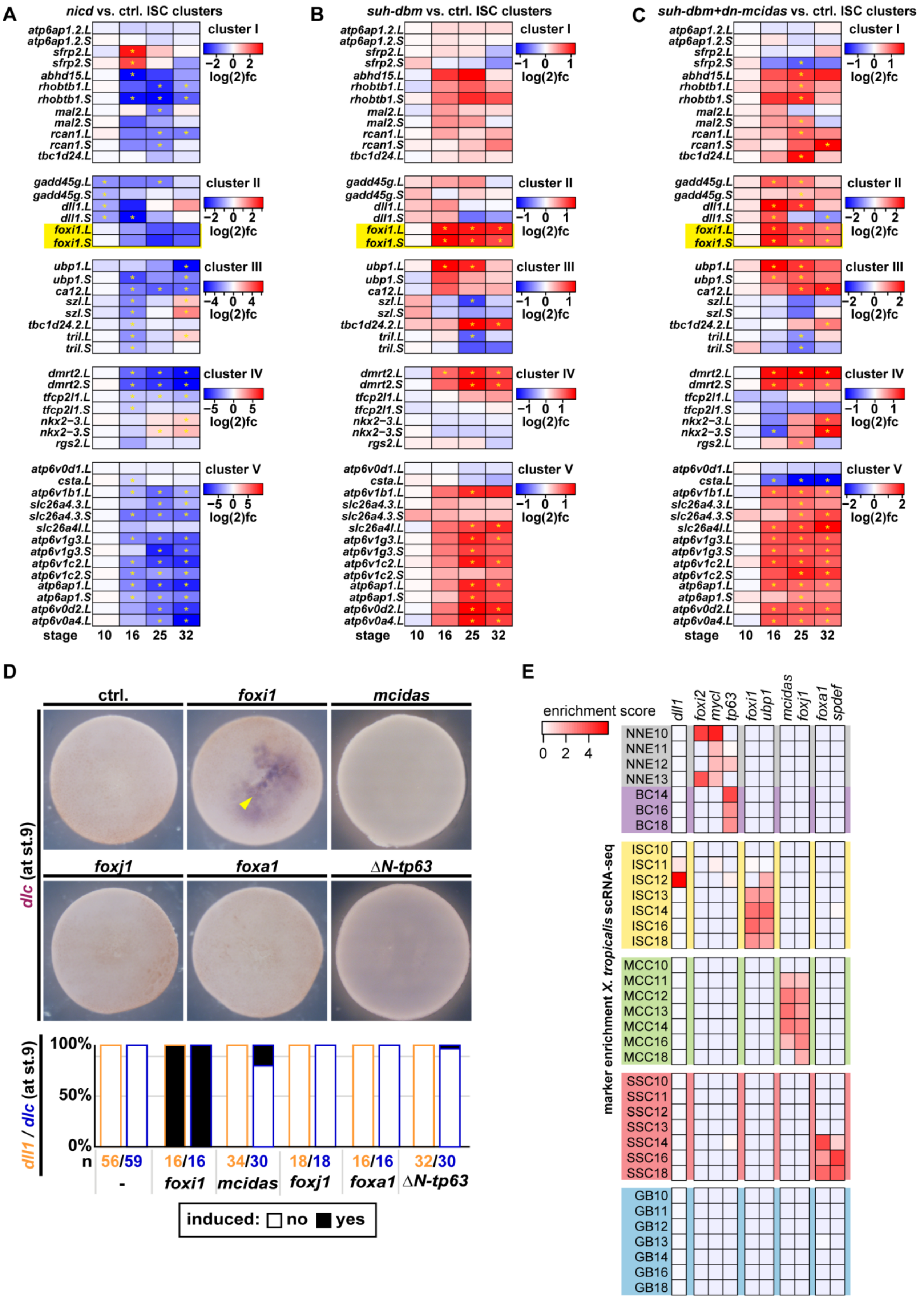

### Figure S7: Notch regulation of ISC genes and ISC-subtype markers

**(A,B,C)** Effects of Notch gain (*nicd*; **A**), Notch loss (*suh-dbm*; **B**) and Notch and MCC loss (*suh-dbm* + *dn-mcidas*; **C**) on core ISC gene expression in key developmental stages (st. 10, 16, 25, 32). Heatmaps depict log2-fold change values derived from DEseq2. Asterisks indicate statistical significant (adj-p value < 0.05) changes. **(D)** Representative images of st. 9 control (ctrl.) and manipulated embryos (animal views) after mRNA overexpression of transcription factors to test premature induction of *dlc*. Quantification of results and effects on *dll1* (yellow) and *dlc* (blue) graphs. Embryos were scored as induced or non-induced expression. Related to Fig. 5A. **(E)** Heatmap of mucociliary marker gene enrichment during differentiation in lineages from scRNA-seq data published in Briggs et al., 2018. Values were derived using the published online tool: kleintools.hms.harvard.edu/tools/currentDatasetsList_xenopus_v2.html. NNE = non-neural ectodermal precursors; BC – basal cells; ISC = ionocytes; MCC = multiciliated cells; SSC = small secretory cells; GB = outer-layer goblet cells.
